# Environment-Dependent Modes of Adaptive Evolution in Neuroblastoma: Plasticity as a Selectable Trait

**DOI:** 10.1101/2023.12.07.570359

**Authors:** Cecilia Roux, Sian Hamer, Abigail Shea, Emilia Chen, Ayeh Sadat Sadr, Christie English, Alejandro Allo Anido, Harvey Che, Birgit Geoerger, Louis Chesler, Gudrun Schleiermacher, Michael David Morgan, Alejandra Bruna

## Abstract

Neuroblastoma, like many aggressive cancers, exhibits phenotypic heterogeneity, contributing to therapy resistance and disease progression. However, direct evidence of how phenotypic plasticity influences tumour evolution remains limited.

Using a multi-resolution quantitative approach, we define the principles, types, and dynamics of plasticity in neuroblastoma at unprecedented resolution. We demonstrate that intrinsic plasticity is a model-dependent process that enables the coexistence of drug-sensitive and drug-resistant states in environmentally stable conditions, positioning plasticity as a bet-hedging strategy that allows tumours to anticipate environmental changes.

Additionally, we show plasticity varies across lineages and single-cell-derived clones, establishing that it is not merely an induced response but a heritable and selectable trait. By simultaneously mapping plasticity at and clonal dynamics in evolving neuroblastoma populations, we show plasticity is shaped by selective pressures, reinforcing its role as a fundamental driver of neuroblastoma evolution in a treatment- and genetic background-dependent manner. We define three distinct modes of plasticity-led adaptation. In the selection of the plasticity model, strong selective pressure temporarily constrains phenotypic transitions, but they reemerge with greater dynamics once the stressor is removed, favouring the selection of highly plastic clones. In the adaptive plasticity model, phenotypic transitions actively reshape tumour heterogeneity, allowing for rapid adaptation to treatment while minimising the impact of clonal selection. Finally, in the plasticity equilibrium model, phenotypic transitions persist at baseline, maintaining a state of phenotypic fluidity, with clonal selection ultimately dictating tumour evolution.

These findings highlight the diverse, context-dependent strategies that neuroblastoma populations employ to navigate selective pressures and therapy resistance, emphasizing the need for plasticity-targeting therapeutic approaches to disrupt tumour adaptation and improve treatment outcomes.

**Teaser:** Natural selection and phenotypic transitions shape adaptive evolution, guiding neuroblastoma survival under treatment pressure.

## Introduction

Paediatric cancers account for more than one in five deaths among children under 14 (Cancer Research UK). Unlike adult cancers, which often arise from an accumulation of somatic mutations, paediatric cancers typically emerge *in utero* due to disrupted differentiation and uncontrolled growth of transformed embryonic cells. These cancers are driven by relatively few genetic alterations, yet they also frequently evade treatment due to their adaptive evolution capabilities. The embryonic cells from which paediatric cancers arise are inherently plastic during development, a property essential for their ability to differentiate into diverse cell types. The developmental origin of childhood cancers, coupled with their low mutational burden, suggests that non-genetic mechanisms, rather than the extensive acquisition of novel mutations, play a key role in treatment resistance and relapse. As such, paediatric cancers offer a unique model for studying plasticity-driven adaptive evolution in cancer.

Neuroblastoma accounts for 8-10% of childhood cancers and 40% of patients survive less than 5 years due to treatment resistance leading to relapse(*1*). Neuroblastoma tumours are usually found in the adrenal gland or sympathetic ganglia and are thought to arise during early embryonic development due to the oncogenic transformation of multipotent trunk neural crest (NC) cells or their immediate derivatives, such as sympathoadrenal/Schwann cell progenitors and sympathoblasts. High-risk disease, which makes up 61% of cases, is frequently characterized by amplification of the oncogenic transcription factor MYCN. Additional recurrent alterations include chromosomal copy number aberrations (e.g. gain of chromosomes 17 (chr17q) and 1 (chr1q)(*2–4*)) and mutations in ALK, ATRX and RAS pathway genes(*2–4*). Comprehensive genomic profiling at diagnosis and relapse has shown evidence of continued or parallel acquisition of structural variants constrained to these driver loci, pointing towards convergent evolution. However, the absence of consistent novel genetic events at relapse suggests these alone do not account for the full complexity of neuroblastoma evolutionary strategies observed in patients(*5*).

Additionally, high-risk neuroblastoma tumours display significant phenotypic heterogeneity(*6–10*), which has been linked to the distorted differentiation trajectories adopted by trunk NC-derived cells(*11*, *12*). An alternative hypothesis proposes that this heterogeneity could instead be driven by developmental plasticity, the ability of a single genotype to generate diverse phenotypes in response to environmental changes.

Single-cell RNA sequencing (scRNA-seq) studies in preclinical and clinical samples have yielded conflicting results, due to the complexity of the neuroblastoma phenotypic landscape and the limitations of the traditional clustering method(*6–10*). While most evidence supports the presence of at least three major cell phenotypic populations reminiscent of different stages of sympathoadrenal development(*6*, *13*), reconciling these findings across patient tumours, patient-derived models, and long-term cultured cell lines remains challenging. Transcriptional programs of neuroblastoma cell states differ between cell lines, patient-derived models, and clinical samples. These discrepancies contribute to apparent inconsistencies in the literature, emphasising the need for a unified framework to systematically study phenotypic heterogeneity and its implications for neuroblastoma evolution and therapy response.

To overcome the limitations of predefined cell state nomenclature and to focus on the dynamics of phenotypic transitions we propose a unifying classification. ADRN (adrenergic) and MES (mesenchymal) cell states, which are defined by distinct core regulatory circuits (CRCs) and CD44 expression levels, are retained as previously established states in neuroblastoma cell line models(*14*). MES is the most extreme non-ADRN state identified to date with transcriptional programs resembling the most undifferentiated stages of the developing sympathetic nervous system. Here, we introduce semi-stable transcriptional states (SSCSs) as a new nomenclature for model-specific transcriptionally defined cell states with unique stability and dynamic properties. This classification accounts for the fluidity of neuroblastoma phenotypic transitions, allowing for a more evolutionarily relevant framework to study plasticity and its interplay with clonal selection in tumour adaptation, and therapy response. Transitions between these cell states, particularly the well-documented ADRN-to-MES transitions (AMTs) observed in preclinical models provide direct evidence of plasticity in neuroblastoma(*14*). However, some studies suggest that MES-like cells only emerge post-treatment(*15*, *16*), while others challenge their presence in primary tumours entirely(*8*, *9*, *17*), raising questions about their role in disease progression and therapeutic resistance.

Here, we address key knowledge gaps regarding plasticity-driven adaptive evolution in neuroblastoma by integrating classical experimental evolution, lineage barcoding, scRNA-seq, and computational modelling. Our findings reveal that neuroblastoma is not merely an induced response but rather a predetermined, heterogeneous, and selectable trait shaped by the environment during adaptation to treatment.

Consistent with natural evolution, our results reveal a complex interplay between clonal selection and phenotypic transitions, which is highly environment-dependent. Briefly, we identified three major plasticity-related modes of adaptive evolution in neuroblastoma preclinical models. 1/ A “selection of plasticity” model, where phenotypic transitions are temporally constrained under strong selective pressure but re-emerge at higher dynamics once the stressor is removed, this model is driven by the selection of highly plastic clones. 2/ An “adaptive plasticity” model, where phenotypic transitions actively reshape tumour heterogeneity, enabling rapid and efficient adaptation to treatment while minimizing the impact of clonal selection. 3/ A “plasticity equilibrium” model, where phenotypic transitions remain active at baseline, maintaining a state of phenotypic fluidity that facilitates ongoing adaptation, ultimately governed by clonal selection. Together, these findings highlight the diverse strategies by which neuroblastoma populations navigate selective pressures, underscoring the context-dependent nature of plasticity in neuroblastoma evolution and therapy resistance.

Understanding how plasticity evolves, shapes tumour heterogeneity, and interplays with clonal selection to facilitate survival under treatment will be crucial for developing targeted therapeutic strategies that disrupt plasticity-driven adaptation and prevent resistance.

## Results

### Three-dimensional phenotypic landscapes reveal intrinsic neuroblastoma plasticity dynamics

Evidence suggest phenotypic heterogeneity is a key determinant of cancer evolution and is well established in both preclinical and clinical models of neuroblastoma(*6–10*, *17*, *18*). However, conflicting findings have been yielded regarding the origins of this heterogeneity, and its evolutionary consequences remain unclear. This has prompted further investigation into the role of plasticity in response to treatment.

To characterise phenotypic heterogeneity in neuroblastoma, we examined preclinical models, including three established cell lines and three patient-derived models of distinct genomic features relevant to neuroblastoma (table 1). We confirmed phenotypic heterogeneity in all models tested based on clustering methods applied to scRNA-seq data (Supplementary Figure 1).

To begin exploring the role of plasticity in neuroblastoma phenotypic heterogeneity at population level, we focused our initial experimental approaches on the well-characterized, plastic SK-N-SH cell line. This widely used model exists in the prototypical neuroblastoma cell states, ADRN and MES. ADRN cells are governed by core regulatory circuits (CRCs) involving transcription factors such as PHOX2B, GATA3, and HAND2, which drive neuronal differentiation programs, whereas MES cells are regulated by AP-1, PRRX1, and RUNX1, promoting a mesenchymal-like, migratory, and therapy-resistant phenotype. These previously identified ADRN and MES states can be readily detected in SK-N-SH cells based on CD44 expression levels, providing a significant advantage for isolating, quantifying, and functionally studying distinct neuroblastoma phenotypes(*14*). To this end, we employed CD44-based fluorescence-activated cell (FACS) sorting (Figure 1a) to derive populations of cell states designated CD44^low^-ADRN, CD44^int^-INT and CD44^high^-MES thereafter (Figure 1a and Supplementary Figure 2a). Quantification of these cell states at different time points after sorting revealed that all three populations exhibited distinct proportions of the other phenotypes, indicating plasticity (Figure 1b). Notably, CD44^int^-INT populations displayed a tenfold higher transition rate to CD44^high^-MES by day 2, while CD44^low^-ADRN populations transitioned rapidly to CD44^int^-INT and CD44^high^-MES states, with CD44^low^-ADRN cells comprising ∼50% of the population by day 12 (Figure 1b). In contrast, CD44^high^-MES populations showed slower transitions, maintaining an 80% CD44^high^-MES composition by day 12. These results indicate that while all neuroblastoma cells retain plastic capabilities independently of their original cell state, the rate and direction of transitions vary in a cell state-specific manner.

**Figure 1.**
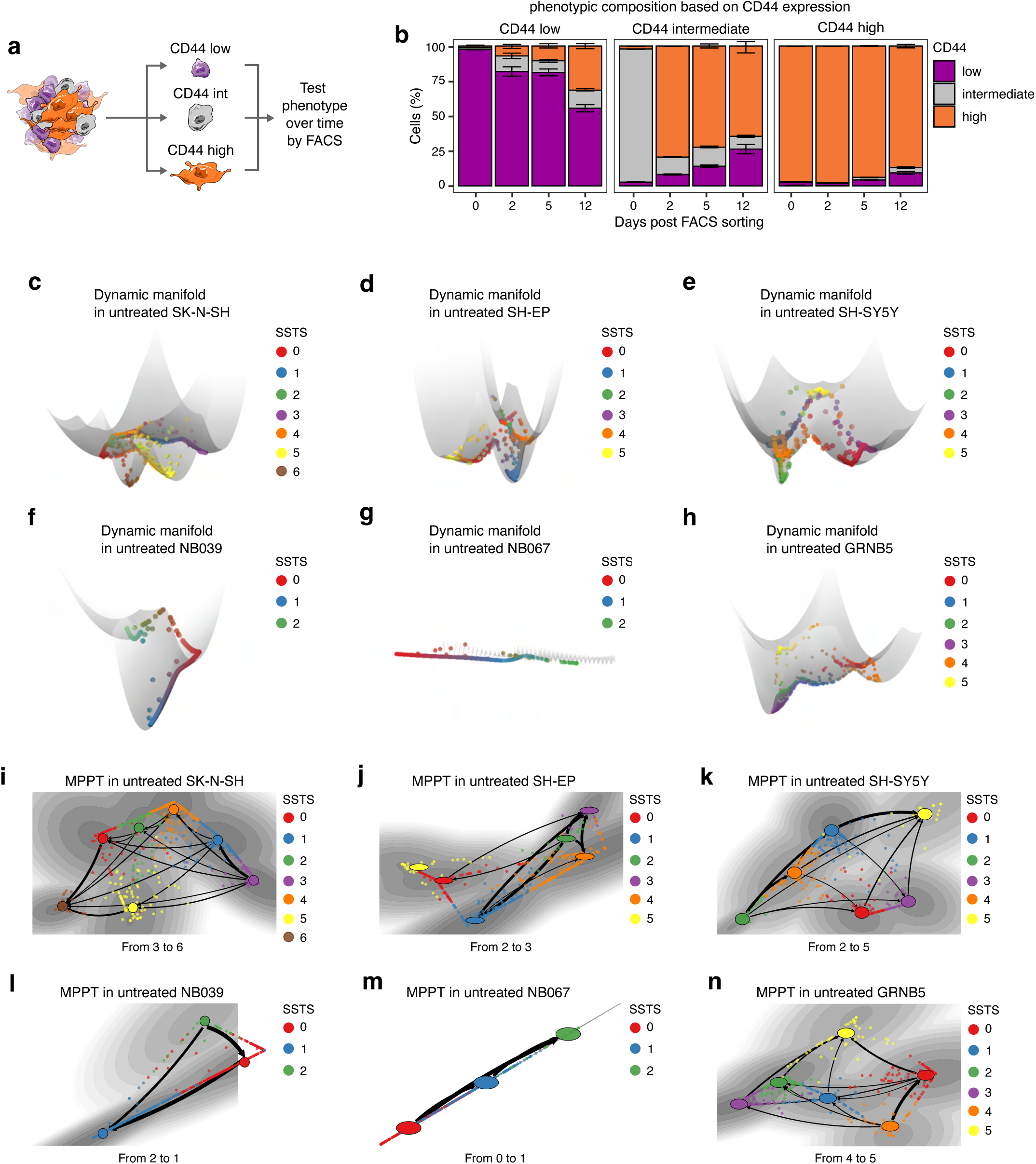
Three-dimensional phenotypic landscapes reveal neuroblastoma plasticity dynamics. **a**, SK-N-SH cells were FACS sorted based on CD44 expression level into three categories (low, intermediate and high) and cultured separately. The phenotypes of these sorted cells were assessed at 2, 5, and 12 days after the FACS sorting using Flow Cytometry. **b**, Stacked bar charts illustrating the percentage of live cells in each CD44 expression category at different time points post-FACS sorting. Error bars display standard deviation (SD) of three technical replicates per time point of a representative experiment. **c-h**, Dynamic manifold plots in treatment naïve SK-N-SH (**c**), SH-EP (**d**), SH-SY5Y (**e**), NB039 (**f**), NB067 (**g**) and GRNB5 (**h**) neuroblastoma cell lines and patient-derived models. **i-n**, Most probable path trees (MPPT) in treatment naïve SK-N-SH (**i**), SH-EP (**j**), SH-SY5Y (**k**), NB039 (**l**), NB067 (**m**) and GRNB5 (**n**) neuroblastoma cell lines and patient-derived models. Both the dynamic manyfold and the MPPT plots are coloured by attractor wells.

To further characterise these phenotypic states, we examined morphological and functional differences. CD44^low^-ADRN cells exhibited higher proliferation rates, CD44^int^-INT cells displayed the highest colony-forming ability, and CD44^high^-MES cells were more resistant to standard chemotherapies, consistent with prior observations (Supplementary Figure 2 and Supplementary Figure 3)(*14*, *19*). These results suggest diverse cell phenotypes exist in neuroblastoma populations each with unique cellular and functional properties. Notably, the presence of cell states with enhanced drug tolerance supports the notion that plasticity functions as an evolutionary bet-hedging strategy, enabling neuroblastoma populations to transition between cellular phenotypes and maintain the coexistence of pre-adapted states capable of surviving therapeutic pressures.

To investigate the existence of phenotypic transitions in naïve, unsorted neuroblastoma populations, we exploited the scRNA-seq-derived transcriptomes across the six models (Supplementary Figure 1), applying both 2D and 3D computational methods. First, we assigned pseudotime ordering to cells and mapped them onto a phenotypic continuum(*20*) (Supplementary Figure 4a-f). Pseudotime and transition scores from extreme phenotypes in both cell lines and patient-derived models were positively correlated, particularly when distinct phenotypes were present (Supplementary Figure 4g-l), supporting the notion that plasticity involves continuous transcriptional changes rather than discrete switches.

To further refine our understanding of plasticity dynamics, we applied dynamical manifold approaches, using the multiscale method MuTrans to construct three-dimensional phenotypic landscapes(*21*). This analysis revealed a model-specific phenotypic topology and demonstrating greater cell state heterogeneity than previously anticipated (Figure 1c-h).

Each cell state was represented as an attractor well, where depth reflected stability: deeper wells indicated more stable states requiring higher energy to transition out of, whereas shallower wells corresponded to more plastic, transient states. Annotation of these attractor wells based on transcriptional profiles (Supplementary Figure 5a-f) confirmed the presence of ADRN and ADRN-like states, alongside other extreme and intermediate profiles that did not fully align with the previously defined ADRN and MES states in the SK-N-SH cell line. This granular resolution analysis highlights the existence of tumour-specific semi-stable transcriptomic cell states (SSTS), providing a refined framework to study plasticity dynamics.

Notably, we observed plasticity in both SH-EP and SH-SY5Y cell lines, which are clones derived from SK-N-SH cells traditionally considered MES- and ADRN-fixed respectively. This challenges the assumption that these clones exist in static, fixed states and suggests that plasticity may be a broader feature of neuroblastoma cell populations. Additionally, phenotypic landscapes differed significantly between the three patient-derived models. The two *MYCN*-amplified neuroblastoma models, NB039 and NB067, exhibited a similar number of attractor states yet differed dramatically in their phenotypic landscape topology. NB039, with wild-type *ALK*, contained one highly stable SSTS among its three attractor states, whereas all SSTS in NB067, with mutant *ALK*, were highly unstable and plastic. In contrast, neuroblastoma cells in the also *MYCN*-amplified patient-derived model GRNB5 exhibited a greater complexity in the types and dynamics of phenotypic transitions and stability patterns, further highlighting model-specific plasticity dynamics and stability landscapes.

To determine the most probable phenotypic transition pathways, we mapped the global transition structure between SSTSs using maximum probability flow trees (MPFT) (Supplementary Figure 5g). We then inferred transition probabilities using the most probable path trees (MPPT) approach (Figure 1i-n). The width of the connecting lines indicated the likelihood of each transition, with thicker lines representing higher probability pathways. Notably, in all models, transitions were predominantly routed through highly plastic (less stable) cell states (i.e. shallow attractor wells). These findings highlight the biological significance of plastic, transient states in facilitating transitions between more stable states.

Overall, these results broaden our understanding of neuroblastoma phenotypic diversity, revealing a wider spectrum of phenotypic states in preclinical models than previously recognized and demonstrating that transitions between these states occur along a continuum. This supports plasticity as a fundamental feature of neuroblastoma, enabling cells to navigate complex phenotypic landscapes shaped by functionally distinct SSTSs with unique topologies and transition dynamics.

### Single-cell derived cultures demonstrate plasticity is a selectable trait in neuroblastoma

To determine whether plasticity-driven phenotypic transitions act as a heritable trait rather than a purely inducible response, we adapted Luria-Delbrück fluctuation principles, incorporating principles of evolutionary reaction norms to quantify plasticity as a trait(*22*).

We established multiple independent single-cell-derived cultures from CD44^low^-ADRN, CD44^int^-INT and CD44^high^-MES from SK-N-SH parental cells. These populations were then expanded in culture for multiple generations before analysing their phenotypic composition based on the proportion of cells in each phenotypic state.

If plasticity is a selectable trait, we would expect fluctuations in phenotypic composition across cultures, indicative of lineage-specific heterogeneity in plasticity. Conversely, if plasticity is purely environmentally induced, we would expect low variance between cultures, suggesting it is an induced response to environmental cues (Figure 2a). Using flow cytometry to classify cells into cell states based on CD44 expression, ten weeks after single-cell isolation revealed a wide spectrum of phenotypic composition, consistent with plasticity being a selectable trait (Figure 2b). The diversity of plasticity patterns observed ranged from varying degrees of phenotypic transitions, leading to different proportions of CD44^low^-ADRN and CD44^high^-MES cells within the population, to low-frequency irreversible transitions culminating in the emergence of CD44^high^-MES-only cultures. Notably, CD44^high^-MES-derived single cells failed to establish viable cultures, suggesting MES states may require pre-existing cellular networks or supportive signals to sustain long-term growth.

**Figure 2.**
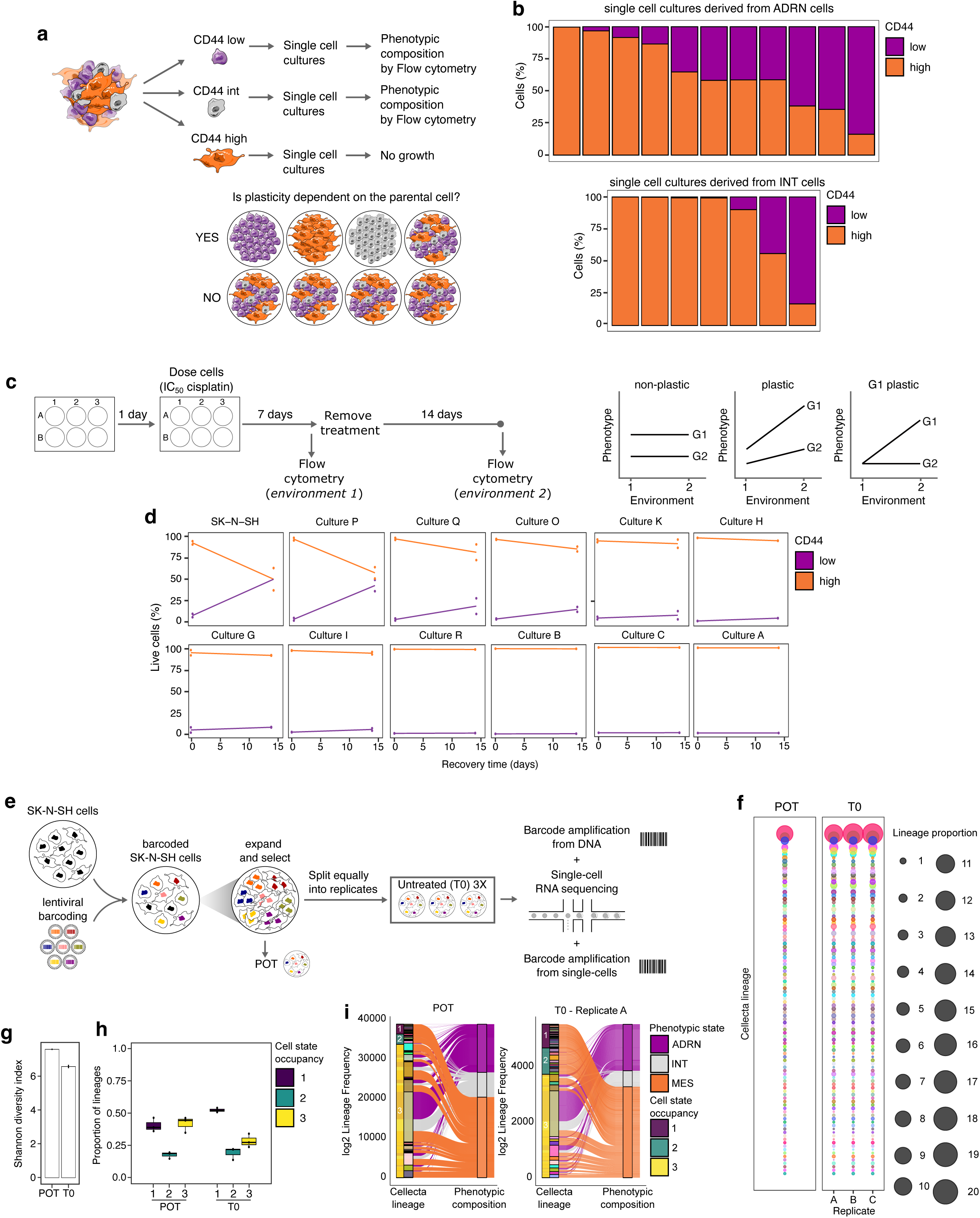
Plasticity diversity amongst neuroblastoma lineages. **a**, Schematic representation of the experimental design and possible experimental outcomes to determine the heritability of plasticity traits. SK-N-SH cells were sorted by FACS into CD44-low, CD44-intermediate and CD44-high populations. Single cell clones were generated using limited dilution approaches and the phenotypic composition of clones were determined by Flow Cytometry. **b**, Stacked bar chart displays the phenotypic composition of single cell clones after approximately 10 weeks in culture, determined by flow cytometry. The clones where derived from either pure ADRN cells (top) or INT cells (bottom). Live cells were gated into CD44-low and CD44-high populations and plotted as the percentage of CD44-low and CD44-high (y axis) for each single cell clone (x axis). Clones were ordered based on the percentage CD44-high cells on this first analysis. **c**, Schematic representation of the reaction norms experimental design (left) and graphical outcomes for plasticity traits (right). G1 and G2 refer to alternative heritable traits. **d**, Reaction norms graphs for parental SK-N-SH cells, and single cell-derived clones from pure ADRN cells. Cells were analysed by flow cytometry immediately after cisplatin treatment (day 0; environment 1) and after 14 days recovery (day 14; environment 2). Live cells were gated into CD44-low and CD44-high populations and plotted as the percentage of CD44-low and CD44-high cells (y axis) at each time point (x axis). Two replicates were analysed per time point. **e**, SK-N-SH cells were tagged with Cellecta barcode library in a lentiviral system and expanded. The founder population (referred to as POT) was divided equally into replicates. The untreated cells were collected and cryopreserved for further DNA and single cell analysis when they reached confluency. **f**, Frequency of the 100 most highly abundant lineages within POT visualized across untreated samples, ordered most to least frequent. These lineages were then visualised across experimental conditions using a dot plot, where size reflects percentage representation of each barcode within a sample, where each colour is associated with a unique lineage. **g**, Shannon diversity index calculated across all lineages per experimental condition visualised as a bar plot; bars display the mean Shannon index score across replicates. **h**, Box plot representing the proportion of lineages associated to either one unique cell state, two cell states or three cell states in POT and untreated samples. **i**, Alluvial plot depicting cell states associated with each Cellecta lineage in POT and a representative untreated sample. Lineages are categorized by the number of cell states, and the thickness of the bar indicates their frequency in the population. First bar coloured by proportion of lineages associated to cell states, second bar coloured by unique lineage, third bar coloured by phenotypic state. Alluvial plot are coloured by phenotypic state.

Importantly, the phenotypic composition of the single-cell-derived cultures remained stable over time and across multiple passages, as confirmed by longitudinal tracking (Supplementary Figure 6). This stability over generations further supports the notion that plasticity is not solely an inducible response but a heritable, lineage-specific trait.

To expand from these analyses and further validate pre-existing heterogeneity in plasticity trait, we quantified plasticity employing a reaction norms approach in a dynamic preclinical setting of drug responses (Figure 2c). Rooted in evolutionary biology, reaction norms describe how a phenotype (y) responds to environmental changes (x), with the slope of the reaction norm quantifying the degree of plasticity(*23*, *24*). We assessed reaction norms within a treatment framework, exposing eleven CD44^low^-ADRN derived single-cell-derived cultures (Supplementary Figure 7a) and parental SK-N-SH cells to cisplatin for 7 days (environment 1), followed by 14 days of drug withdrawal in drug-free media (environment 2). These conditions were chosen based on observations that environment 1 led to a CD44^high^-MES-only culture in parental SK-N-SH cells, which, upon drug withdrawal (environment 2), repopulates the original phenotypic structure indicating plasticity(*25*) (Figure 2d). By determining the proportion of phenotypes in each environment we observed that parental SK-N-SH yielded a reaction norm slope of 3.0 (Figure 2d, Supplementary Figure 7b). Single-cell-derived cultures displayed varying degrees of responses, reflecting a spectrum of plasticity traits. Cultures P, Q, and O showed plasticity reverting to mixed ADRN/MES populations in environment 2, with reaction norm slopes of 2.8, 1.1, and 0.8, respectively. In contrast, cultures A, B, and C maintained a pure MES state with a slope of 0.0 despite arising from an ADRN single cell, demonstrating a lack of plasticity capability. Using this evolutionary method that provides a quantitative framework for measuring plasticity, we demonstrate that plasticity exhibits varying strengths across individual lineages within a neuroblastoma population, aligning with the definition of a heterogeneous trait.

We next investigated whether this plasticity inherent to the cell of origin influences the overall behaviour of the population. To address this, we analysed single-cell-derived cultures and found that lineage-specific plasticity positively correlated with growth rates and treatment sensitivities to cisplatin (Supplementary Figure 7c-f). Cultures derived with low or no plasticity capability exhibited slower proliferation and reduced cisplatin sensitivity. In contrast, highly plastic single-cell derived cultures displayed faster growth and increased cisplatin sensitivity (Supplementary Figure 3e,f and Supplementary Figure 7b-f).

### Plasticity diversity at single-cell resolution in naïve populations using DNA barcoding

To further investigate plasticity at single-cell resolution in naïve, unperturbed populations, we applied a DNA-barcoding approach (Figure 2e). These barcodes could be detected in both DNA and RNA sequencing, enabling simultaneous lineage tracing and transcriptomic profiling allowing for high-resolution lineage tracing, enabling us to track cell state transitions over time while maintaining information on clonal identity. By leveraging DNA barcodes, we can investigate whether plasticity-driven phenotypic transitions occur stochastically across all cells or if they are lineage-dependent, providing further evidence of its variability and heritability within naïve neuroblastoma populations.

We integrated DNA barcodes into the genome of individual cells, with each barcode uniquely marking a single lineage.

To ensure robust tracking of cell state transitions across lineages, the barcode library transduction was optimized to track 100,000 lineages, followed by an 850× library expansion step to increase the likelihood of obtaining identical barcodes across independent replicates (herein sister replicates). However, this initial expansion step led to a loss of diversity, reducing the initial 100,000 potential barcodes to an average of 16,124 unique barcodes in the pool of cells (POT) for further investigations.

We next analysed lineage dynamics by examining the frequency distribution of the top 100 barcodes in cultured cells at T0 (baseline, before treatment) across sister replicates. The Shannon index was also calculated for all lineages to assess lineage diversity and stability over time (Figure 2f,g). Lineage frequency distributions and diversity indices were highly consistent across all untreated conditions, both in sister replicates (T0s) and in longitudinal samples (POT and T0s), indicating that pre-existing lineages consistently survived and proliferated under culture conditions.

To assign quantitative language to plasticity at the lineage level, we integrated lineage and transcriptomic data and assigned plasticity scores based on the proportion of cell states (defined by transcriptional programs) per barcode (Figure 2h,I and Supplementary Figure 8a). We observed that approximately 50% of the lineages in the naïve population exhibit some degree of plasticity. For example, in the POT sample, 40.5% of lineages were linked to a single phenotypic state, 17.6% to two states, and 41.9% to three states, indicating that phenotypic transitions occurred in most of the population. Additionally, we observed distinct plasticity patterns. Lineages exhibiting one or two phenotypic states were predominantly MES and/or INT, whereas ADRN cells were more commonly found coexisting with multiple other cell states (Figure 2i and Figure 1b). A linear correlation between lineage frequency and the number of states occupied was further observed (Supplementary Figure 8b).

Studying plasticity using approaches at a single-phenotype population level (adaptation to Luria-Delbrück fluctuation assays) and at single-cell lineage (DNA-barcoding) resolution, our findings provide strong evidence that phenotypic transitions continuously occur in naïve populations in all phenotypic states and in multiple directions, albeit at different rates and with distinct transition dynamics.

### Treatment-specific adaptive strategies interplay with neuroblastoma phenotypic heterogeneity

Having established a quantitative framework to quantify plasticity and confirmed that plasticity is a cell-of-origin trait, we next investigated whether and how plasticity interplays with clonal selection to drive adaptive evolution in response to treatment. To address this, we extended the dynamic drug response framework, which features ON/OFF treatment cycles, previously used in the reaction norm approach (Figure 2c). This mimics the key phases of therapy-induced adaptation, capturing the transition into a drug-tolerant persister (DTP) or persistence phase, followed by awakening and regrowth. We selected clinically and preclinically relevant compounds, including chemotherapeutic agents and epigenetic modifiers (epi-drugs) (Supplementary Figure 9a), and performed this analysis on the SK-N-SH cell line. The proportion of CD44^low^-ADRN, CD44^int^-INT and CD44^high^-MES cells was determined at different time points of the drug response process revealing treatment-specific shifts in cell state composition. To further investigate neuroblastoma evolutionary strategies in response to changes in the environment, we selected three prototypical compounds representative of the main phenotypic dynamics observed in response to treatment: cisplatin (which led to enrichment for CD44^high^-MES consistent with environment 1 in the reaction norms, Figure 2d), the EZH2 inhibitor (EZH2i) Tazemetostat (which led to enrichment for CD44^low^-ADRN), and the BRD4 inhibitor (BRD4i) JQ1 (which led to no significant changes on the phenotypic composition).

Previous studies suggest cisplatin-mediated enrichment on MES-like cells is due to induced plasticity, where highly proliferative ADRN cells transition to resistant phenotypes to increase the chances of survival to treatment(*25*). However, without temporal resolution, it remains unclear whether plasticity is the primary adaptation mechanism or if cisplatin simply exerts strong selection against ADRN cells, leaving pre-existing MES cells to persist and survive treatment. To investigate this, we examined the survival of isolated CD44^low^-ADRN and CD44^high^-MES cells in response to cisplatin. We observed a complete elimination of the CD44^low^-ADRN-sorted population, whereas some CD44^high^-MES cells persisted in cisplatin treatment (Figure 3a,b and Supplementary Figure 3e). Notably, cisplatin-persistent CD44^high^-MES cells displayed increased resistance to subsequent cisplatin exposure compared to untreated CD44^high^-MES cells (Figure 3c, left panel). Upon treatment withdrawal, surviving CD44^high^-MES cells awakened and regrew, restoring the original phenotypic population structure (Figure 3b) and regaining cisplatin sensitivity (Figure 3c, right panel), indicating the surviving CD44^high^-MES-DTPs are highly plastic and capable of replenishing proliferative ADRN-like states.

**Figure 3.**
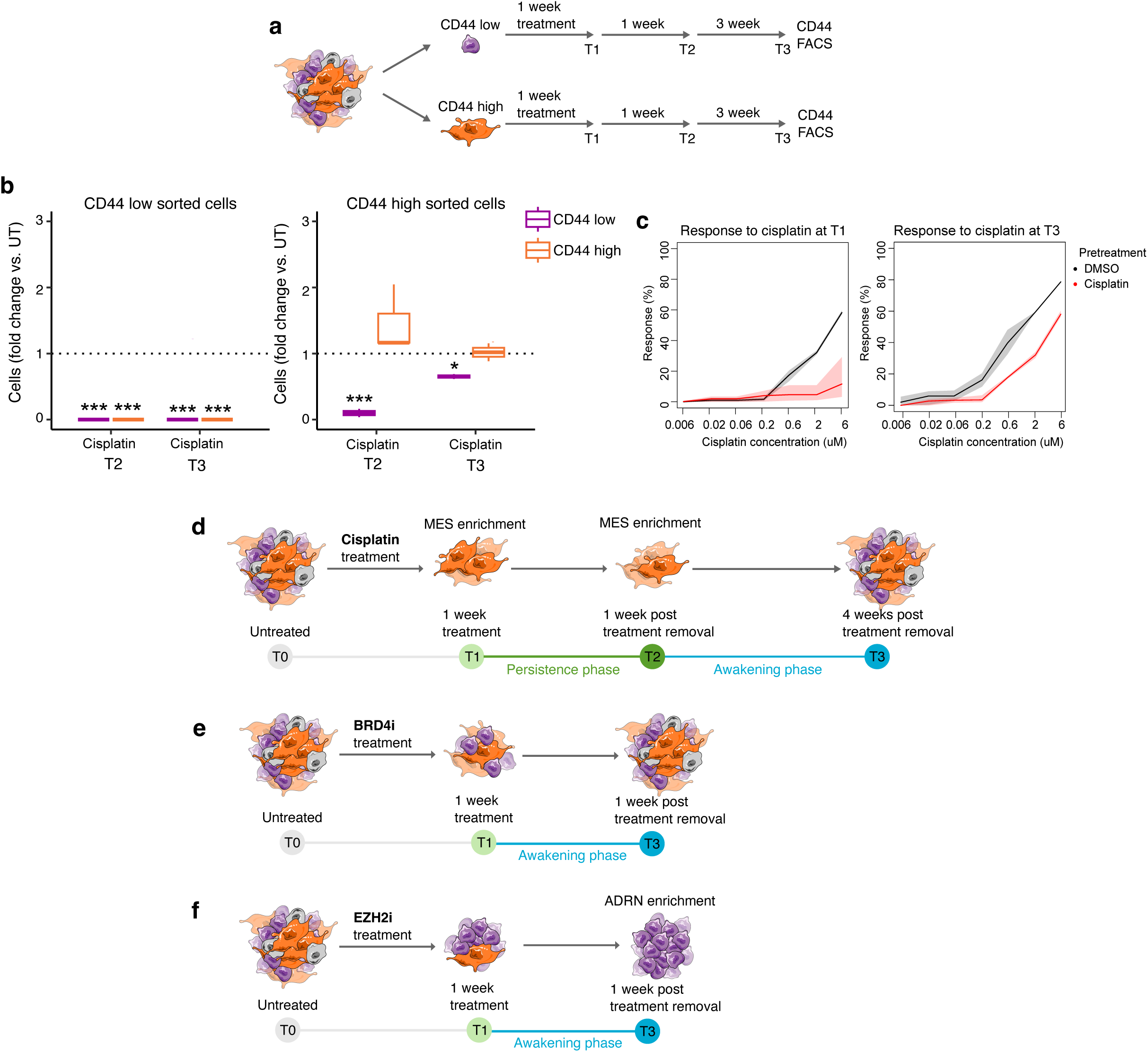
Divergent treatment-specific adaptive strategies interplay with neuroblastoma phenotypic heterogeneity. **a**, SK-N-SH cells were FACS sorted based on CD44 expression and each population was treated with cisplatin 1 week. Following the removal of treatment, CD44 expression was evaluated by Flow Cytometry after 1 week (T2) and 4 weeks (T3). **b**, Box plots displaying the CD44-low and CD44-high populations within each experimental condition. Values are expressed as fold change relative to the untreated samples and plots display mean ± SD of biological replicates for a representative experiment. Statistical significance is denoted using asterisks (*P ≤ 0.05, **P ≤ 0.01, ***P ≤ 0.001) based on t-test comparing the results to the untreated control. **c**, Cisplatin drug-response curves of SK-N-SH cells collected at T1 (left panel) and T3 (right panel) time points after cisplatin treatment. For each plot, untreated SK-N-SH cells were used as controls. The response rate is represented as percentage and shaded area displays confidence intervals. **d-f**, Schematic of treatment framework to model the drug-induced entry into a persistence phase (named T1 and T2) and the subsequent awakening of the cells after long term treatment removal (named T3) for cisplatin (**d**), BRD4i (**e**) and EZH2i (**f**).

Treatment with EZH2i led to an enrichment of CD44^low^-ADRN cells without evidence of selective killing, as the working concentration was below the cytotoxic threshold (Supplementary Figure 3e). This suggests that plasticity was the primary mechanism of adaptation, consistent with previous findings (Supplementary Figure 8e)(*26*).

In contrast, BRD4i treatment did not induce detectable changes in cell state composition, despite significantly reducing cell viability. This indicates that selection is prioritised while plasticity was not engaged as a relevant adaptive strategy under these conditions (Supplementary Figure 8b).

These findings highlight how different therapeutic agents exploit or bypass neuroblastoma plasticity to drive drug tolerance, persistence and regrowth. Cisplatin selects for plastic lineages, EZH2i primarily drives phenotypic reprogramming and BRD4i induces cell death independently of plasticity, suggesting that its effects are not mediated by phenotypic transitions (Figure 3d-f).

### Treatment- and patient-specific modulation of the phenotypic landscape

To validate and expand on from these observations, we performed scRNA-seq analysis at multiple time points of the drug response process (Figure 3d-f), leveraging the collection of cell lines and patient-derived models described in Figure 1. scRNA-seq data showed treatment-specific transcriptomic changes (Supplementary Figure 10, Supplementary Figure 11 and Supplementary Figure 12). We probed the transcriptomic data using pseudotime and dynamic manifold approaches as described in Figure 1. Pseudotime analysis revealed multiple trajectories within the inferred global lineage structure, indicative of treatment-specific plasticity (Supplementary Figure 10 and Supplementary Figure 13).

Dynamic manifold approaches revealed the phenotypic landscape of each model, characterized by the type and stability of SSTS and their most frequent transition paths, was shaped in a treatment-dependent manner (Figure 4 and Figure 5).

**Figure 4.**
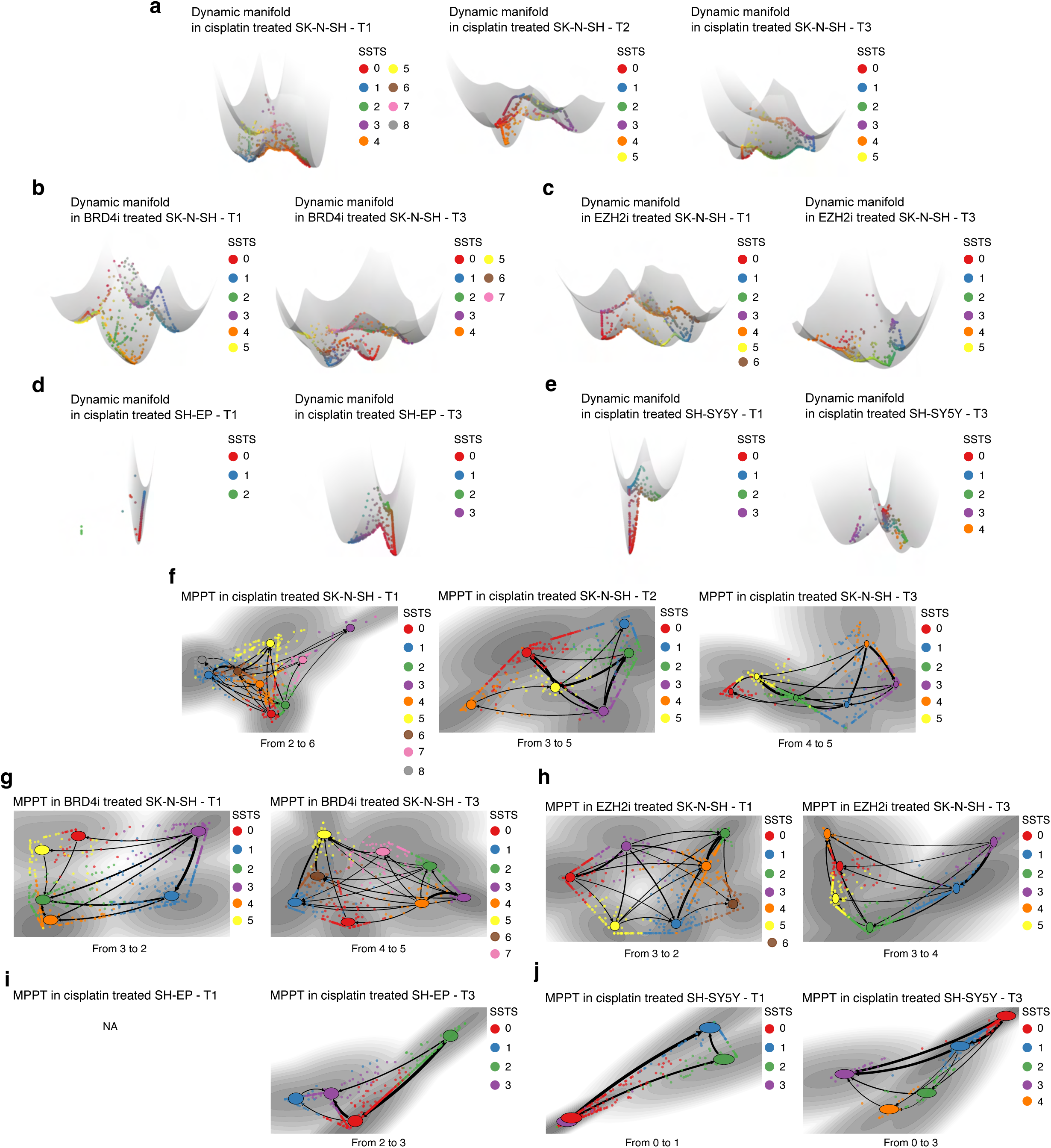
Treatment-specific modulation of the phenotypic landscape in neuroblastoma cell lines. **a-e,** Dynamic manifold plots in cisplatin treated SK-N-SH (**a**), BRD4i treated SK-N-SH (**b**), EZH2i treated SK-N-SH (**c**), cisplatin treated SH-EP (**d**), cisplatin treated SH-SY5Y (**e**) neuroblastoma cell lines. **f-j**, Most probable path trees (MPPT) in cisplatin treated SK-N-SH (**f**), BRD4i treated SK-N-SH (**g**), EZH2i treated SK-N-SH (**h**), cisplatin treated SH-EP (**i**), cisplatin treated SH-SY5Y (**j**) neuroblastoma cell lines. Both the dynamic manyfold and the MPPT plots are coloured by attractor wells and divided by experimental time point. The thickness of the arrows in the MPPT plots is reflective of the probability of transition. The MPPT plot for SH-EP could not be computed due to an insufficient number of cells.

**Figure 5.**
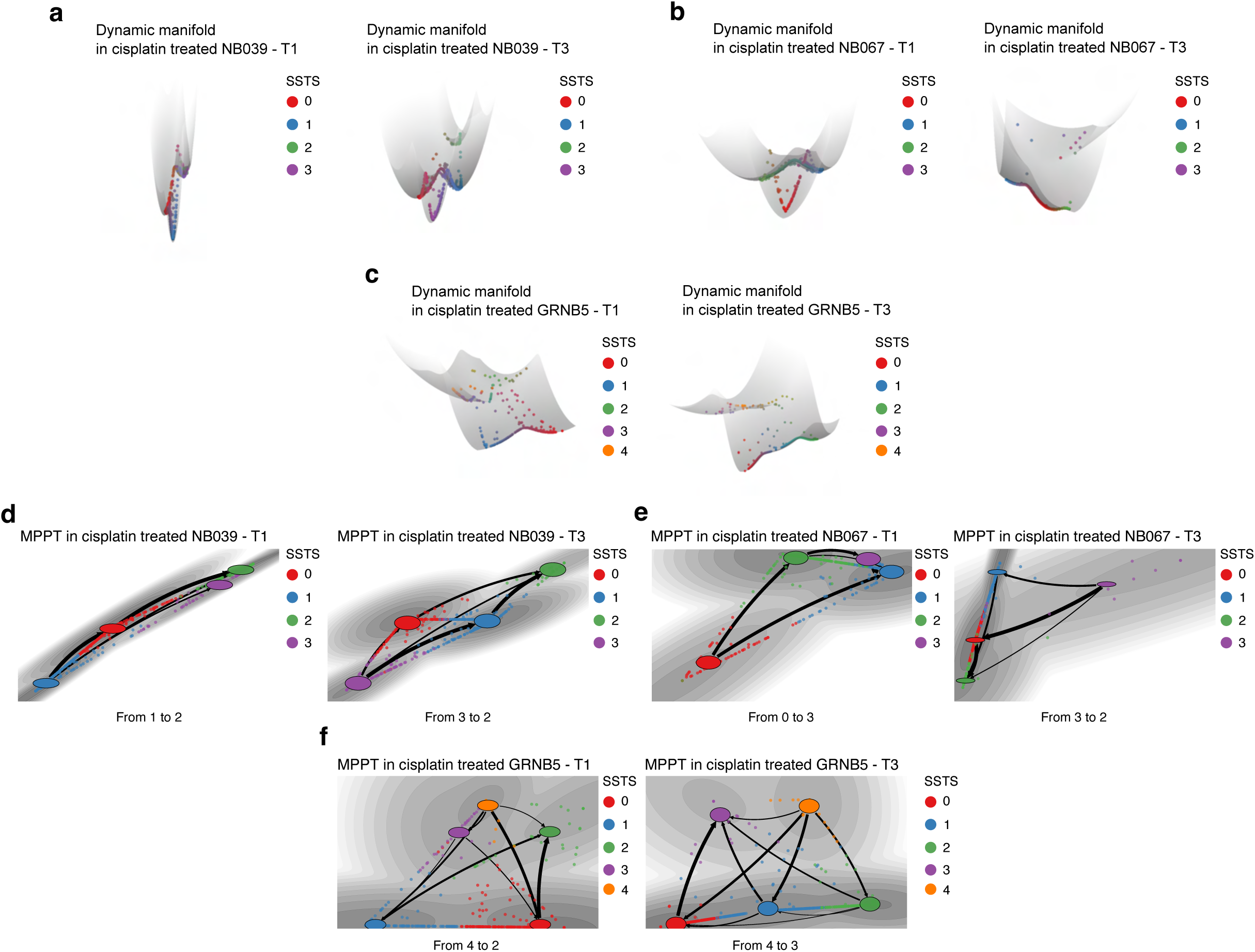
Treatment-specific modulation of the phenotypic landscape in neuroblastoma patient-derived models. **a-c**, Dynamic manifold plots in cisplatin treated NB039 (**a**), cisplatin treated NB039 (**b**), cisplatin treated GRNB5 (**c**) neuroblastoma patient-derived models. **d-f**, Most probable path trees (MPPT) in cisplatin treated NB039 (**d**), cisplatin treated NB039 (**e**), cisplatin treated GRNB5 (**f**) neuroblastoma patient-derived models. Both the dynamic manyfold and the MPPT plots are coloured by attractor wells. The thickness of the arrows in the MPPT plots is reflective of the probability of transition.

In SK-N-SH cells, cisplatin treatment increased the stability of the few ADRN-like SSTSs while decreasing the stability of multiple other SSTSs, compared to untreated controls. This increase in unstable attractor wells suggests that cells explore a broader phenotypic space, likely in search of fitter states for survival. Upon awakening (T3), the phenotypic landscape reverted to a topology resembling the original population, indicating that plasticity-driven adaptation is reversible following drug withdrawal (Figure 4a,f and Supplementary Figure 14).

EZH2i treatment induced subtle topological alterations, yet the phenotypic landscape exhibited a global decrease in SSTS stability, suggesting that EZH2i promotes an increase in phenotypic flexibility, potentially enabling cells to transition more readily between states (Figure 4c,h and Supplementary Figure 15). In contrast, BRD4i treatment restricted phenotypic transitions across all SSTSs simultaneously, as indicated by the increased stability of most attractor wells (Figure 4b,g and Supplementary Figure 15). This global stabilisation of SSTSs under BRD4i treatment likely explains why previous analyses failed to detect major shifts in phenotypic composition (Supplementary Figure 9b). These findings demonstrate that static analyses alone may overlook key plasticity-driven adaptation mechanisms, reinforcing the necessity of dynamic resolution in studying plasticity.

The SK-N-SH clones, SH-SY5Y and SH-EP, exhibited distinct phenotypic landscape shifts in response to cisplatin treatment. In the ADRN-enriched SH-SY5Y cell line, cisplatin treatment induced a transition towards a single predominant stable SSTS, suggesting a strong constraint on phenotypic transitions. However, upon drug withdrawal, the post-treated culture recovered multiple SSTSs with varying stability levels, mirroring the effects observed in SK-N-SH parental cells. This reversion to the original, untreated, phenotypic landscape indicates that plasticity remains an accessible mechanism for adaptation following awakening after a strong environmental pressure (Figure 4e,j and Supplementary Figure 16). In contrast, the MES-enriched SH-EP clone, which lacks the Ink4 locus, exhibited resistance to phenotypic landscape changes in response to cisplatin (Figure 4d,i and Supplementary Figure 16). This suggests that its phenotypic stability may be inherently canalised by genetic constraints, limiting plasticity-driven transitions and reinforcing a genetically imposed restriction on adaptation dynamics.

In the MYCN-amplified, ALK-mutant patient-derived model (NB067), cisplatin exposure dramatically altered the phenotypic landscape topology, shifting from a highly plastic, flat topology to a constrained and stable landscape dominated by a single SSTS. Upon drug withdrawal, the landscape partly reverted to its original highly plastic topology, indicating that plasticity was temporarily suppressed but not permanently lost (Figure 5a,d and Supplementary Figure 17). In contrast, the MYCN-amplified, ALK wild-type models NB039 and GRBN5 did not exhibit major topological shifts in response to cisplatin, but phenotypic transitions were observed continuously throughout the drug response process (Figure 5b,c,e,f and Supplementary Figure 17 and Supplementary Figure 18). This suggests that rather than experiencing changes in plasticity, these models maintain a dynamic equilibrium, where phenotypic transitions persist throughout treatment adaptation and regrowth, enabling continuous evolutionary flexibility.

### Clonal and plasticity dynamics to refine distinct neuroblastoma adaptive strategies

To increase the granularity of our observations and refine our understanding of the relationship between lineage and transcriptomic dynamics, we applied previously defined barcoding approaches within the treatment framework on the SK-N-SH cell line at single-cell resolution.

We first analysed lineage dynamics during the treatment process (Figure 6) to determine whether pre-existing fitter clones influence clonal behaviour. If such clones existed before treatment, we would expect similar clonal frequencies and distributions across sister replicates (Figure 6a). Indeed, we observed consistent clonal dynamics in sister replicates across most conditions, with the notable exception of awakening replicates following cisplatin withdrawal (T3).

**Figure 6.**
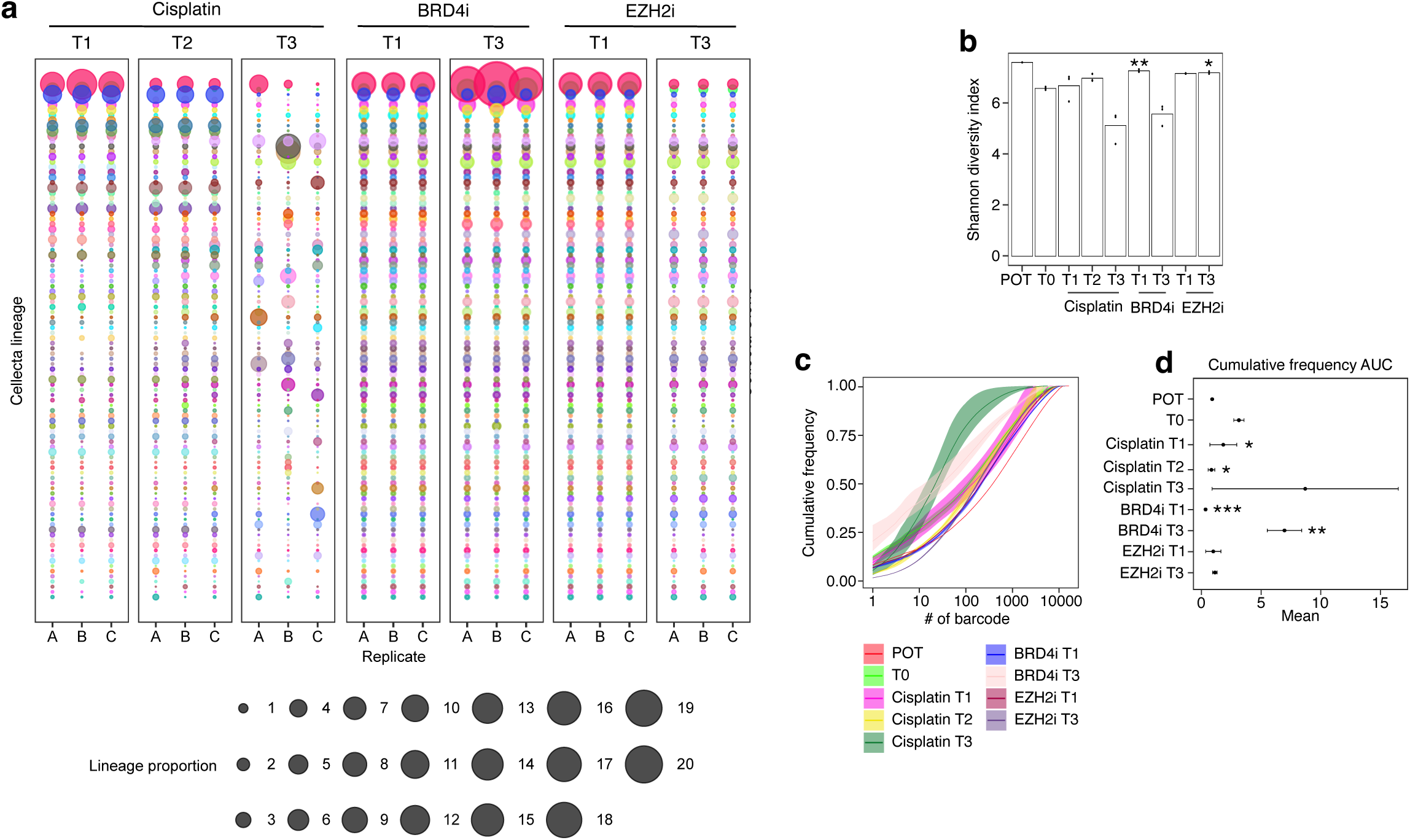
Selective bottlenecks during treatment responses. **a**, Frequency of the 100 most highly abundant lineages within POT (see Figure 2f) visualized across all other samples and replicates, ordered most to least frequent. These lineages were then visualised across experimental conditions using a dot plot, where size reflects percentage representation of each barcode within a sample, where each colour is associated with a unique lineage. **b**, Shannon diversity index calculated per experimental condition visualised as a bar plot; bars display the mean Shannon index score across replicates. **c**, Cumulative proportion distribution plots calculated per experimental condition (lines and variance coloured by experimental condition), visualising cumulative barcode/lineage ranked from most to least frequent. **d**, AUC relative to the cumulative proportion distribution plot in **c**. **e**, Frequency of the 100 most highly abundant lineages within each awakened (T3) sample, visualized across the previous time points and replicate, ordered most to least frequent. These lineages were then visualised across experimental conditions using a dot plot, where size reflects proportion representation of each barcode within a sample, where each colour is associated with a unique lineage. Colour scheme for unique lineages is common across all experimental arm plots.

Shannon diversity index calculations and cumulative frequency analysis revealed that cisplatin imposed the strongest selection bottleneck, indicating a selective sweep of fewer highly fit clones dominate (tolerate and persist treatment and then regrow) (Figure 6b-d). BRD4i treatment also induced selection, but dominant clones are accumulating more slowly, suggesting that the awakened post-treatment population is more evenly distributed across different lineages (Figure 6b-d). For EZH2i treatment, we did not observe a selection advantage despite changes in lineage frequencies, which is consistent with our previous evidence suggesting primarily plasticity-driven adaptation (Figure 6b-d).

We next calculated the proportion and type of cell states within lineages at different time points to track plasticity dynamics throughout the drug response process. During the persister phase, cisplatin treatment significantly reduced plasticity, as evidenced by a decrease in the number of lineages with multiple states (Figure 7a,b). Similarly, plasticity was lost under BRD4i treatment but retained in response to EZH2i (Figure 7c, Figure 6b-d). During awakening, we observed an increase in the proportion of lineages with >2 cell states, compared to untreated cultures, particularly in response to cisplatin and EZH2i consistent with the MuTrans analysis (Figure 7a-c, Supplementary Figures 16a,b).

**Figure 7.**
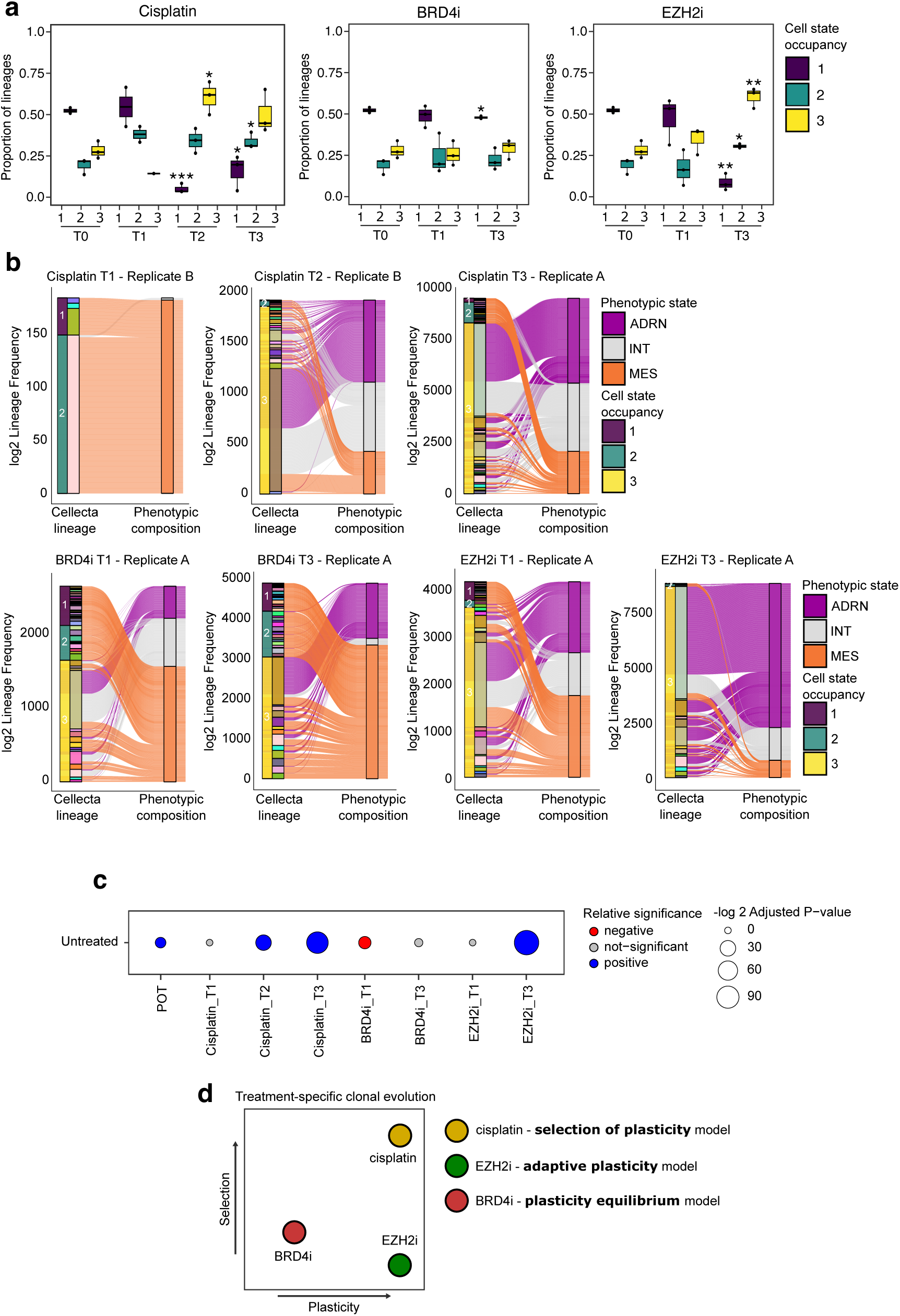
Selection of plastic lineages during treatment responses lead to an increased plastic population. **a**, Box plot representing the proportion of lineages associated to either one unique cell state, two cell states or three cell states in cisplatin, BRD4i or EZH2i treated samples. Plots display mean ± SD of biological replicates for a representative experiment. Statistical significance is denoted using asterisks (*P ≤ 0.05, **P ≤ 0.01, ***P≤0.001) based on comparisons between each experimental condition and untreated samples. Statistical significance was determined using pairwise t-tests and Bonferroni multiple-comparison correction methods. Where no asterisks are found on plot there was no significant difference between comparison after Bonferroni multiple comparison correction. **b**, Alluvial plot depicting cell states associated with each Cellecta lineage representative treated sample. Lineages are categorized by the number of cell states, and the thickness of the bar indicates their frequency in the population. First bar coloured by proportion of lineages associated to cell states, second bar coloured by unique lineage, third bar coloured by phenotypic state. Alluvial coloured by phenotypic state. **c**, Dot plot illustrating the chi-squared test significance comparing the frequency of cells in 1 or more than 1 cell states for each treatment arm and time point, compared to the untreated sample. The size of the dot represents the -log2 adjusted p value and the colour indicates either a non-significant comparison (grey), greater proportion of cells in > 1 cell state than untreated (blue) or fewer proportion of cells > 1 cell state than untreated (red). **d**, Schematic illustration of the treatment-specific clonal evolution modes influenced by selection and plasticity.

The selection of plasticity at awakening post cisplatin and EZH2i treatment was further accompanied by treatment-specific shifts in phenotypic transition patterns (Figure 7b). This highlights that plasticity-driven phenotypic transition types and dynamics change during drug responses, reinforcing the context-dependent nature of plasticity in adaptive evolution.

Altogether, our findings suggest at least three distinct plasticity-dependent adaptive strategies exist in neuroblastoma populations: 1/ selection of plasticity, where treatment selects for highly plastic clones but temporarily constrains it to enable survival of drug resistance cells; 2/adaptive plasticity, where induced phenotypic transitions actively shape tumour heterogeneity in response to the environment, enabling rapid adaptation to treatment while limiting clonal selection; and 3/plasticity equilibrium, where continuous phenotypic transitions maintain a fluid phenotypic landscape, facilitating ongoing phenotypic flexibility through clonal selection.

Together, these findings highlight the diverse, context-dependent strategies that neuroblastoma populations employ to survive therapeutic pressure.

### Investigating the impact of plasticity evolution on the population fitness

To further explore the adaptive strategy involving the selection of plasticity, we focused on cisplatin treatment, given its clinical relevance as a frontline chemotherapeutic agent. Having established that plasticity evolves in response to cisplatin treatment, we next aimed to determine how this evolution influences key evolutionary parameters, such as fitness. We hypothesised that highly plastic dominant lineages that undergo natural selection will enhance the overall fitness of the evolving population.

To test this, we expanded upon classical evolutionary frameworks (Figure 2c,d) and quantified plasticity changes across multiple rounds of cisplatin treatment and awakening (Figure 8a).

**Figure 8.**
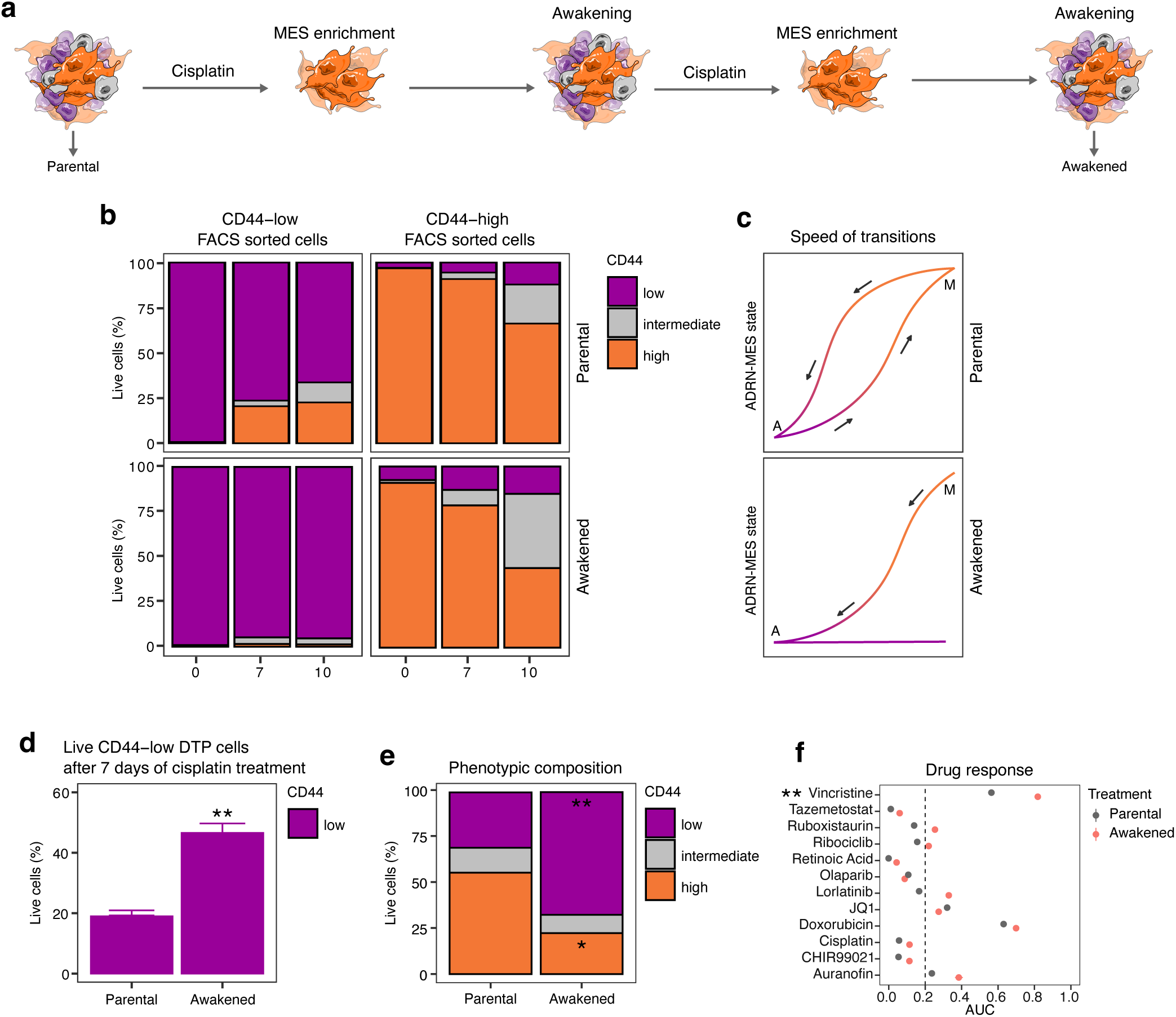
Plasticity changes during treatment drives opposing cell state adaptation to enhance functional fitness. **a**, Schematic illustration of the experimental framework. SK-N-SH cell were treated with cisplatin for 7 days, following which treatment was removed for 4 weeks. The same schedule of treatment was administered twice, the parental and awakened cells were analysed to assess the plasticity capacity of the population. **b**, Stacked bar charts illustrating the percentage of live cells in each CD44 expression category at different time points post-FACS sorting in either parental of awakened population. Error bars display standard deviation (SD) of three technical replicates per time point of a representative experiment. **c**, Schematic illustration of the speed of cell state transitions in parental and awakened populations. **d**, Percentage of live CD44-low cells 7 days after cisplatin treatment in parental and awakened populations. **e**, Stacked bar chart illustrating the percentage of live cells gated base on CD44 expression into low, intermediate and high, in parental and awakened populations. **f**, Drug response profiles of parental and awakened cells. Dot plots display area under the curve (AUC) values. Statistical significance is denoted using asterisks (*P ≤ 0.05, **P ≤ 0.01) based on comparisons between parental and awakened samples. Statistical significance was determined using pairwise t-tests and Bonferroni multiple-comparison correction methods. Where no asterisks are found on plot there was no significant difference between comparison after Bonferroni multiple comparison correction.

Using this approach, we monitored and quantified cell phenotypes, DTPs, drug responses, and plasticity dynamics across multiple cisplatin treatment cycles in SK-N-SH cells. The results revealed a consistent pattern of cell state-specific plasticity adaptations. Under cisplatin selection pressure, CD44^low^-ADRN cells transitioned from a plastic to a plastic-low state (Figure 8b,c). This loss of plasticity was accompanied by increased cisplatin resistance (Figure 8d), a shift in phenotypic composition at the population level (Figure 8e), and altered drug responses to a panel of clinically relevant compounds (Figure 8f). The transition from a highly plastic, treatment-sensitive state to a plastic-low, treatment-resistant state ensures the survival and evolution of ADRN cells, which were initially preferentially eliminated by the drug.

In stark contrast, CD44^high^-MES cells, which are inherently more resistant to treatment, exhibited a progressive increase in plasticity, enhancing their ability to transition back to the CD44^low^-ADRN state (Figure 8b,c). This increase in MES plasticity expanded the reservoir of cells capable of differentiating into ADRN-like proliferative cells, effectively replenishing the proliferative pool. This aligns with the enrichment of ADRN populations following extended ON/OFF cisplatin treatment cycles, compared to untreated cultures.

This combined adaptive strategy underscores the complex interplay between cell states and plasticity in response to selection pressures, demonstrating that plasticity changes are finely tuned to balance fitness, survival, and functional costs. These findings suggest that evolution optimizes adaptive strategies to maintain tumour heterogeneity and enhance resilience to therapy.

## Discussion

This study provides a comprehensive framework for understanding the role of plasticity in the early stages of neuroblastoma evolution. By integrating experimental evolution, single-cell approaches, and computational modelling, we demonstrate that plasticity is both an induced process and a heterogeneous and selectable trait that dynamically evolves during early adaptive responses to treatment. Our findings emphasise the need to incorporate plasticity dynamics into studies of cancer evolution and translational research, towards developing anti-plasticity therapeutic strategies.

Childhood cancers, such as neuroblastoma, are characterized by disruptions in developmental processes(*11*, *12*), allowing tumours to exploit embryonic mechanisms such as cell plasticity to adapt without requiring additional genetic mutations. This supports the hypothesis that plasticity, rather than genetic selection alone, is an early driver of treatment adaptation, enabling the emergence of high-fitness traits without requiring additional mutations. However, direct evidence linking plasticity to cancer evolution has been limited due to technological constraints. Given that plasticity is only evolutionarily relevant if it facilitates transitions to adaptive, fitter phenotypes, new approaches are essential to capture and quantify plasticity within evolving tumour populations.

By examining plasticity at three complementary levels (isolated cell states, population-level heterogeneity, and single-cell lineage dynamics), in preclinical cell lines and patient-derived models with distinct genetic backgrounds, we demonstrate that neuroblastoma phenotypic heterogeneity in naïve untreated populations is a consequence of plasticity. We show neuroblastoma cells navigate a transcriptional continuum within a phenotypic landscape composed of semi-stable transcriptionally regulated states (SSTSs) with distinct transition dynamics and stability levels. Our analyses uncover previously uncharacterised phenotypic states that expand beyond the known ADRN and MES populations, highlighting that the neuroblastoma phenotypic landscape is influenced by genetic factors. Although the topology of phenotypic landscapes varies across models, phenotypic transitions in all models predominantly route through highly plastic intermediate states rather than through direct shifts between extreme phenotypes. This highlights the biological importance of unstable, highly plastic phenotypes as critical intermediates in neuroblastoma adaptation. The low frequency of these highly plastic cell states strongly suggests that plasticity comes at an evolutionary cost, potentially constraining tumour growth and proliferation as the highest proliferative cells where those with and ADRN-like phenotype. This opens the door to evolutionary vulnerabilities, where targeting plasticity itself, rather than specific cell states, may provide a therapeutic advantage.

By applying classical experimental evolution and lineage barcoding, we further demonstrate that plasticity is not uniformly distributed across populations but rather varies between lineages. These findings challenge the traditional view of neuroblastoma evolution as merely a process induced by treatment(*5*, *27*). Instead, our evidence supports a model where plasticity itself is a trait. In naïve, untreated cultures, some lineages exhibit high plasticity, transitioning between multiple states without external stimuli, whereas others are constrained to specific phenotypic fates. Additionally, the nature and dynamics of phenotypic transitions vary between lineages, reinforcing the idea that plasticity, like genetic mutations, is heritable and subject to evolutionary pressures.

To further investigate whether plasticity is subject to selection, shaped and shaping the trajectory of neuroblastoma evolution, we deepened our studies using three therapeutic agents (cisplatin, EZH2i, and BRD4i) that induce distinct phenotypic dynamics under ON/OFF exposure cycles. At the clonal space, we observed that cisplatin exerted the strongest selection pressure, preferentially selecting for low-frequency pre-existing subclones, leading to a significant reduction in post-treatment population diversity. BRD4i also reduced population diversity, but instead of favouring rare clones, it selected more abundant pre-existing lineages. EZH2i, in contrast, did not significantly alter lineage diversity, suggesting a lower impact of natural selection and a stronger reliance on plasticity-driven adaptation.

To determine whether the selected subclones were pre-adapted, we analysed lineage distributions across independent sister replicates. Lineage composition remained highly consistent across all treatment time points, except in cisplatin awakening, where replicates significantly diverged. This suggests that clonal selection under treatment follows Darwinian principles, where pre-existing subclones best suited to the new environment are preferentially selected. However, the stochastic nature of lineage distributions post-cisplatin withdrawal suggests that following a strong evolutionary bottleneck, the regrowth trajectories of surviving lineages become inherently unpredictable. This reflects the probabilistic nature of awakening and expansion under sparse population densities.

In response to cisplatin, despite clear evidence of selection, we found no transcriptional signatures that could define or predict future clonal dominance. Given our earlier findings that plasticity is a heritable trait, we hypothesised that highly plastic lineages are preferentially selected during treatment. To test this, we used expressed barcode information to trace the transcriptome of lineages that became DTPs, survived, awakened and regrew the post-treated population in all conditions. Our results reveal at least three distinct plasticity-driven adaptive strategies in neuroblastoma evolution. Plasticity selection occurs when the plasticity process is temporarily constrained, but plasticity as a trait is selected, favouring the survival of plasticity-equipped clones that can endure treatment. Plasticity-driven adaptation involves induced plasticity actively remodelling phenotypic heterogeneity, enabling rapid adaptation while minimising clonal selection. In contrast, plasticity equilibrium maintains phenotypic flexibility, allowing for ongoing transitions while selection prioritises other fitness traits. These distinct strategies highlight the context-dependent role of plasticity in shaping neuroblastoma adaptation, emphasising the need to consider both plasticity dynamics and selection pressures when designing therapeutic interventions.

In response to cisplatin, we observed model-specific plasticity landscape alterations in preclinical models. Although no single genetic determinant correlated with these differences, our results suggest that plasticity is influenced by the genetic background of the tumour population. Future studies incorporating a larger number of models will be essential for identifying molecular regulators of plasticity-led evolution.

Altogether, our evidence suggests a model where phenotypic plasticity plays a fundamental role in shaping neuroblastoma evolution, in a complex interplay between natural selection and induced transitions through the phenotypic landscapes in response to environmental pressures. One major discovery of our work is the evidence that plasticity is itself a selectable trait, highlighting its potential as a targeted therapy to block evolution and as a predictive biomarker for relapse risk. This paves the way for future research aimed at mapping plasticity histories and identifying epigenetic signatures associated with plasticity-driven adaptation. Targeting plasticity, rather than individual cell states, may be a more effective strategy to improve treatment outcomes for high-risk neuroblastoma patients. Future research should focus on identifying molecular regulators of plasticity-led adaptation, particularly the epigenetic mechanisms driving plasticity shifts. Additionally, leveraging plasticity-based biomarkers for early relapse detection could revolutionize neuroblastoma treatment by enabling patient stratification and personalized therapeutic interventions targeting plasticity-driven evolution.

## Methods

### Cell lines and treatment

SH-EP, SH-SY5Y and SK-N-SH human neuroblastoma cells were cultured in DMEM (Thermo Fisher) supplemented with 10% fetal bovine serum (FBS) (Thermo Fisher), 100 IU penicillin and 100 μg/ ml streptomycin (Sigma). All cell lines were originally provided by L. Chesler, ICR, London. For SK-N-SH, SH-EP and SH-SY5Y assays compounds were administered at the following concentrations: 3.18 uM cisplatin (Cambridge Bioscience), 0.66 uM BDR4i JQ1 (Selleck Chemicals) and 1 uM EZH2i tazemetostat (Selleck Chemicals).

### Flow cytometry and sorting

For antibody staining of cell surface molecules, isolated cells were aliquoted into 5mL polystyrene FACS tubes, and washed with 2 mL PBS supplemented with 1% BSA (referred to as FB buffer). Cells were pelleted at 400 g for 5 minutes at 4 °C, resuspended in 100 μL FB, and incubated for 30 minutes on ice with the following antibody: anti-CD44 APC/Fire 750 (1:100, Biolegend, 103061). Stained cells were washed twice by centrifugation in 2 ml FB at 400g for 5 minutes at 4°C. Washed cells were resuspended in 25 ng/mL DAPI (Miltenyi), and analysed by flow cytometry (Symphony A5, BD Biosciences) or sorted by FACS (Symphony S6, BD Biosciences; SONY MA900 or SONY SH800, SONY) as above. The data were analysed with FlowJo software.

### CFSE-based proliferation assay

Cells were labelled with 0.5uM CFSE (Invitrogen) following manufacturer’s instructions and incubated for a maximum of 12 days. CFSE dilution was measured by flow cytometry. The data were analysed with FlowJo software.

### Immunofluorescence

For immunofluorescence staining, 2× 10^5 cells were seeded in a 24-well plate containing glass coverslips. The day after, cells were treated with either Cisplatin or JQ1 for the indicated amount of time. After treatment, cells were fixed with 4% PFA for 30 minutes at 4°C. Cells were then permeabilized with 0.2% Triton X-100 for 10 minutes at RT. Cells were washed twice and incubated in PBS buffer containing 1% BSA and 2% FCS (IFF buffer) for 30 minutes at room temperature (RT). Cells were stained overnight at 4 °C with primary antibody diluted in IFF buffer (anti-CD44 IM7, 1:500, eBioscience; anti-LGR5 OTI2A2, 1:200, Abcam; anti-GATA3 D13C9, 1:1000, Cell Signalling, anti-gH2A.XpPhospho (Ser139) 2F3, 1:200, Biolegend). The following day, cells were washed three times with PBS and subjected to conjugated secondary antibody diluted in IFF buffer (anti-Mouse-AF488 A21202 Invitrogen, anti-Rabbit-568 A11036 Invitrogen, anti-Rat-AF647 A78947 Invitrogen) at 1:2000 for 2h at RT. Coverslips were mounted with DAPI Fluoromount-G mounting medium and imaged the with a confocal microscope (Zeiss LSM700).

### Dose response curves

Cell lines were seeded at 2000 cells/well in 384-well plates and cultured for 24 hours prior to dosing. Cells were dosed on the Echo 550 liquid handler, using a 7-dose response curve per compound with three technical replicates per concentration. Cell viability was assessed using CellTiterGlo-3D (Promega) at day 3 after dosing. Cell viability was normalised to background and solvent controls, and dose response curves were fitted using isotonic regression, as previously described(*28*).

### Colony forming assay

For the colony forming assay cells were seeded at 1000 cells/well of a 6-well plate in complete media. Colony growth was monitored, and media was replenished once a week. Plates were fixed and stained with methanol and Giemsa once colonies reached sufficient size to be counted with the naked eye.

### Single-cell derived cultures generation

SK-N-SH cells were sorted into CD44-low (ADRN), CD44-int (INT) and CD44-high (MES) populations by FACS. Sorted cells were seeded in 12-well plates (∼50,000 cells/well) and cultured for 4 days. Cells were then reseeded in 96-well plates to generate single-cell cultures using limiting dilution approaches (seeding density of 0.5 cells/well), in 50% conditioned media and 50% fresh culture media. Fresh media was added to cells once per week, and each single cell clone was expanded for approximately 10 weeks. Single cell clones were then analysed by flow cytometry (as described above) to determine the proportion of ADRN and MES cells.

To generate conditioned media, SK-N-SH cells were seeded at low density and cultured for 5 days without refreshing the culture media. After 5 days, media was collected from culture flasks and centrifuged at 250xg for 10 minutes. The top 90% of media was collected and filtered (0.22um) prior to use.

### Reaction norms and functional characterisation of single cell derived clones

Single cell clones from pure ADRN cells and parental SK-N-SH cells were seeded at 200,000 cells/well in 2 mL of media in 6-well plates using 4 replicate wells per sample. 24 hr after plating, cells were dosed with cisplatin (3.18 uM). 7 days after dosing, 2 wells per sample were harvested and analysed by flow cytometry for CD44 expression (as described above). The remaining 2 wells per sample were washed once with PBS and 2 mL of fresh media was added. 14 days after drug withdrawal, 2 wells per sample were harvested and analysed by flow cytometry for CD44 expression (as described above). Fresh media was added once per week during the drug withdrawal phase.

Cisplatin drug responses and growth dynamics of the single cell clones and parental SK-N-SH cells were analysed in parallel with reaction norms. Cisplatin drug response was tested in two ways: dose response curve and response to a fixed concentration. The dose response curves were generated using the protocol described above using a 6-dose range of 2, 0.6, 0.2, 0.06, 0.02, 0.006 uM and cell viability was measured after 3 days using CTG-3D (Promega). To test the response to a fixed concentration of cisplatin 20,000 cells were seeded in a 24-well plate using 3 replicates per condition. 24 hr after plating, cells were dosed with 3.18 uM cisplatin (to match reaction norms) or left untreated. Cell viability was measured 7 days after dosing using resazurin (final concentration 25 ug/mL, Abcam).

To assess growth dynamics, cells were seeded at 4000 cells/well in 200 uL of media in 96-well plates using 5 replicate wells per sample. Images were acquired using the Incucyte S3 at 10x magnification every 6 hours for 7 days. Percentage confluency was calculated based on surface area coverage using Incucyte (Sartorius) image analysis software.

### Barcoding of cell lines

The CloneTracker library was a gift from Andrea Sottoriva (BCXP1M3RP-XSP, Cellecta). SK-N-SH cell lines were cultured in normal growth media and barcoded by lentiviral transduction according to manufacturer’s recommendations. Following lentiviral titration, 1 million neuroblastoma cells were transduced with a lentiviral library using a multiplicity of infection (MOI) of 0.1 (corresponding to 10% of cells transduced) to attain a single barcode per cell. 100,000 barcoded cells were FACS-sorted based on RFP-positivity and expanded. This population constitutes the “original population” or POT.

### Experimental design of single cell RNA sequencing of barcoded SK-N-SH cell line

Barcoded SK-N-SH cells were seeded equally into 4 treatment groups and divided across three replicates. The seeding density was adjusted based on each treatment and a total of 5M cells were seeded for each sample. Untreated cells were collected at confluency. Three samples were exposed to cisplatin treatment (3.18 uM), three to the BRD4i JQ1 (0.66 uM) and three to the EZH2i tazemetostat (1 uM). After 1 week (T1), the treatment was removed and cells were detached, 10% of the total population was cryopreserved in four aliquots of 2.5% and the remaining cells were reseeded at a fixed density and grown under normal growth media for 1 week (T2) or 4 weeks (T3). All the BRD4i and EZH2i treatment groups were collected at 1 week (T3), whereas the cisplatin treatment groups were collected at both time points (T2 and T3). Cell counts were determined via the Countess II Automatic Cell Counter (Thermo Fisher).

### Single cell RNA profiling of barcoded SK-N-SH cell line

After thawing, single cells were washed with PBS and were resuspended with a buffer (Ca++/Mg++ free PBS + 0.04% BSA). Cells were tagged with the CMO oligos (10X Genomics) as per manufacturer’s instructions. Samples were multiplexed in groups of 6, each containing 5000 cells/samples for a total of 30000 cells/pool.

Pooled single cell suspensions were loaded on a Chromium Single Cell 3ʹ Chip (10X Genomics) and were run in the Chromium Controller to generate single-cell gel bead-in-emulsions using the 10X genomics 3ʹ Chromium v2.0 platform as per manufacturer’s instructions. Single-cell RNA-seq and CMO libraries were prepared according to the manufacturer’s protocol and the library quality was confirmed with a Bioanalyzer High-Sensitivity DNA Kit (Agilent, 5067-4627). Cellecta barcode libraries were prepared using 25% of the adaptor-ligated cDNA to amplify only the barcoded cDNA product. Barcodes were pre-amplified using the P5-Read1 primer (ACACTCTTTCCCTACACGACGCTCTTCCGATCT) and the P7-Adapter-FBPI Cellecta primer (GTGACTGGAGTTCAGACGTGTGCTCTTCCGATCTCCGACCACCGAACGCAACGCACGCA), following which Illumina indexes were added. Samples were sequenced on an Illumina NovaSeq 6000 according to standard 10X Genomics protocol, aiming for approximately 50,000 reads per cell for gene expression libraries and 1000 reads per cell for CMO/Cellecta barcode libraries.

### Single cell data pre-processing

10x Genomics CellRanger (v6.1.2) was used to process raw sequencing FASTQs, using cellranger count (or multi in the case of samples multiplexed with CellPlex) pipeline to conduct read alignment to human genome reference (GRCh38-2024-A), sample demultiplexing and generate feature-barcode matrix. For PDX models, read alignment was conducted using combined human and mouse genome reference (GRCh38_and_GRCm39-2024-A), to observe any mouse gene expression contamination within the sample (as shown in Supplementary Figure).

EmptyDrops method from DropletUtils (v1.20.0) package(*29*, *30*) was implemented on cellranger output H5 matrix to remove empty droplets with false discovery rate of 0.1%. Data files were merged and then normalised using logNormCounts from scuttle (v1.10.1) package(*31*) for downstream QC and processing.

Cell-level filtering was carried out with the following thresholds: mitochondrial gene content <10%, ribosomal gene content >5%, and UMI counts >1000 for cell lines, gene content <40%, ribosomal gene content >0.3%, and UMI counts > 1000 for PDO models, gene content <30%, ribosomal gene content >0.2%, and UMI counts > 1000 for PDX models. Filtering at the gene-level was implemented to remove genes with fewer than five reads. Downstream normalisation and scaling were performed using Seurat (v4.3.0)(*32*) with the SCTransform function, using the top 1000 variable genes, and regressing out cell cycle and mitochondrial genes. The SCTransform data output was used to perform a principal component analysis (PCA). The batch correction algorithm Harmony (v0.1.1) was applied across CMO multiplexed pools (where appropriate), and the output used for Uniform Manifold Approximation and Projection (UMAP)-based dimensional reduction. Clustering was carried out using the Seurat FindClusters function and gene expression markers for the clusters were identified by a Wilcoxon Rank Sum test, using the Seurat FindAllMarkers function. Gene set enrichment analysis using clusterProfiler (v4.6.2)(*33*, *34*) was then carried out on the cluster markers. To compare changes in cell clusters, statistical testing was conducted using student t-test between each sample and T0 (untreated), with adjusted p-value calculated using Bonferroni method for multiple-comparison correction.

### AMT score generation

Single-cell expression signatures for the ADRN and MES cell states were created using the AddModuleScore function in Seurat and published gene sets(*15*). To calculate the AMT score we used a method based on that used by Tan et al.,(*35*). Statistical testing (t-tests) was used to determine whether differences between ADRN and MES gene set expression were significant. The t-statistic was taken as the AMT score and the statistical significance was further used to segregate cells into phenotypic groups: ADRN (AMT score < 0; P < 0.05), intermediate ADRN (AMT score < 0; P ≥ 0.05), intermediate MES (AMT score > 0; P ≥ 0.05) and MES (AMT score > 0; P < 0.05). The two intermediate classifications were grouped together as INT.

For the additional models, cells were assigned three cell state scores (p-MES, p-SYMP, or p-ADRN) based on a patient-derived gene signature(*13*). The highest output of AddModuleScore for the three signatures was used to assign cell states.

To compare changes in phenotypic cell state assignment, statistical testing was conducted using student t-test with adjusted p-value calculated using Bonferroni method for multiple-comparison correction.

### Trajectory inference

Trajectory inference was carried out using Slingshot(*21*) (v2.6.0), which was run with the dimensionality reduction produced by PCA, cluster labels identified by Seurat FindClusters (resolution = 0.4 for barcoded SK-N-SH; resolution = 0.4 for additional models) and setting the root to an ADRN cluster. Gene dynamics analysis by fitting a linear spline model to each lineage was carried out using TSCAN (v.1.36.0).

### MuTrans analysis

Pre-processing of the scRNA-seq data, split by drug condition and experimental timepoint, was carried out analogous to as outlined above using Scanpy (v1.9.3)(*36*). Using the filtered data as inputs, the count matrices were normalised and log transformed with a pseudo-count of 1. The number of transcripts per cell and the mitochondrial content were regressed out and the data was scaled, prior to PCA. Dimensionality reduction using UMAP and Leiden clustering was carried out. Cluster markers were then found using Wilcoxon rank-sum tests.

A neighbourhood graph was computed for the processed data using scanpy.pp.neighbors (metric = “cosine”). Construction of cell-fate dynamical manifolds and downstream analysis of cell-state transition was then performed using the MUltiscale method for TRANSient cells(*21*).

### Cellecta barcodes amplification and next generation library preparation

SK-N-SH cells were harvested and pelleted at the relevant timepoints. Genomic DNA was isolated using AllPrep kit (Qiagen) according to manufacturer’s instructions and quantified using Qubit (Life Technologies). Amplicon PCR was conducted using 50ng of DNA (or 20ng for samples with lower DNA yield) along with 2x Accuzyme mix (Bioline) to amplify barcodes using the following primer sequences (Forward - ACTGACTGCAGTCTGAGTCTGACAG, Reverse - CTAGCATAGAGTGCGTAGCTCTGCT). After identifying and purifying PCR product (consisting of 80-bp product + 30-bp semi-random barcode), NGS libraries were prepared using NEBnext Ultra II DNA library preparation kit for Illumina (New England Biolabs) and quantified with using a TapeStation (Agilent Genomics). Samples were balanced and pooled together to have final molarity of 4nM, NGS was then performed by ICR Genomics facility (TPU) using NovaSeq SP PE150.

### Cellecta barcodes bioinformatics analysis

FASTQ files from bulk experiments were mapped to a custom index reference generated from white-list (all possible sequences) of Cellecta barcodes, using Burrows-Wheeler aligner (BWA-MEM tool; version 0.7.17)(*37*). Once mapped BAM files were generated using Sequence Alignmnet/Map (SamTools; v1.18)(*38*). Gene counts were quantified using FeatureCounts from Rsubread. Parameters were set to filter out reads with mapping quality less than 30 (minMQS = 30).

To observe lineage dynamics, representative percentage was calculated for each lineage within each sample (frequency of Cellecta barcode / frequency of all Cellecta barcodes × 100).

Shannon diversity index(*39*) of all DNA barcodes from each experimental samples was calculated between conditions to observe changes in lineage diversity as per following equation: H=−∑[(pi )×log(pi )], where pi is proportion of individual lineage within the population. Statistical significance of results was determined using pairwise *t-tests*.

Cumulative proportion quantification of all DNA barcodes from each sample was calculated by ranking all barcodes from most to least abundant (sum of frequency). Cumulative proportion was then calculated as cumulative sum of barcode abundance / sum of abundance. For statistical quantification an area under the curve (AUC) value was calculated for each sample using *trapz()* (from R package prcama) of cumulative proportion and abundance from cumulative proportion plot. Statistical significance of results was determined using pairwise *t-tests* between untreated and each other samples.

For top 100 POT barcode analysis, barcodes in POT sample were ranked most to least frequent and the top 100 most abundant barcodes were extracted. Remaining experimental samples were filtered for these barcodes and visualised.

For top 100 awakening barcode analysis, barcodes in each T3 sample were ranked most to least frequent and the top 100 most abundant barcodes were extracted. Samples from the same experimental treatment arm were filtered for these barcodes and visualised. Common colour scheme for barcodes of interest at T3 was generated across all top 100 barcodes lists from all experimental arms, to allow visualisation of common barcodes across experimental arms.

For scRNA-seq experiments Cellecta barcodes from 10x Chromium libraries were isolated using STAR solo(*40*) with the same white list of barcodes used for BWA for bulk experiments. Where multiple barcodes were observed within a single cell, cells were assigned the barcode with the higher proportion of representation from read counts. To account for sparse data, lineages were filtered to ensure each barcodes were observed at least 3 times within the entire dataset.

For plasticity score analysis, each lineage was given a plasticity score based on the number of cell states occupied by that lineage. Statistical significance of results was determined using pairwise *t-tests* between untreated and each other samples. To assess the relationship between cell frequency and plasticity group assignment we conducted spearman’s correlation coefficient in each condition. P-value represented on each plot corresponds to correlation coefficient significance across all replicates within each condition.

Plasticity transition dynamics analysis was conducted by assigning each cell within each lineage a plasticity score based on the number of cell states occupied; either =1 for a single phenotypic state or > 1 for multiple phenotypic states. For analysis frequency of cells in each group were aggregated across replicates. Chi-squared statistical testing was conducted in a pairwise fashion to observe whether there was an association between sample and plasticity based on frequency of each plasticity group. Statistical test was conducted between untreated and each other sample and subject to multiple comparison correction using Bonferroni’s method. Directionality of statistical significance was calculated as (frequency of cells in >1 plasticity group / total cell frequency in sample) - (frequency of cells in >1 plasticity group from untreated / total cell frequency in untreated). Positive was termed change value > 0 and negative < 0.

### Patient-derived xenograft establishment

PDX were established from neuroblastoma tumour biopsies obtained at diagnosis or at relapse, as described previously, following informed consent of parents or guardians(*41*, *42*). This study was performed in accordance with the recommendations of the European Community (2010/63/UE) for the care and use of laboratory animals. Experimental procedures were approved by the ethics committee of the Institut Curie CEEA-IC #118 (Authorization APAFIS#11206-2017090816044613-v2) in compliance with the international guidelines. The establishment of PDX received approval by the Institut Curie institutional review board OBS170323 CPP ref 3272; n° de dossier 2015-A00464-45. In brief, tumor samples were engrafted subcutaneously in immunocompromised mice. Upon successful establishment (> P2) and upon reaching ethical size, PDX tumors were cryopreserved for banking, morphological and molecular analyses according to procedures described previously.

### Patient-derived models and treatment for scRNA-seq analysis

The PDX line GR-NB5 ((MYCN^AMP^, previously named MAPGR-B25-NB-117(*25*)) was originally provided by Gudrun Schleiermacher (Curie Institute, Paris, France). Cells were obtained from enzymatical dissociation of the PDX and cells were cultures in in low-Glucose GlutaMAX DMEM (Thermo Fisher) supplemented with 25% Ham’s F-12 Nutrient Mix, 2% FBS (Thermo Fisher), B-27 Supplement (50X) (Thermo Fisher), minus vitamin A N-2 Supplement (100X) (Thermo Fisher), 100 U/mL penicillin-streptomycin (Sigma), 20 ng/mL Animal-Free Recombinant Human EGF (Peprotech), 40 ng/mL Recombinant Human FGF-basic (Peprotech), 200 ng/mL Recombinant Human IGF-I (Peprotech), 10 ng/mL Recombinant Humand PDGF-AA (Peprotech) and 10 ng/mL Recombinant Human PDGF-BB (Peprotech).

NB039 (MYCN^AMP^ ALK^WT^) and NB067 (MYCN^AMP^ ALK^R1275Q^) patient-derived spheroids were originally provided by J. Molenaar (Princess Maxima Centrum, Netherlands). PDOs were cultured in low-Glucose GlutaMAX DMEM (Thermo Fisher) supplemented with 25% Ham’s F-12 Nutrient Mix, B-27 Supplement (50X) (Thermo Fisher), minus vitamin A N-2 Supplement (100X) (Thermo Fisher), 100 U/mL penicillin-streptomycin (Sigma), 20 ng/mL Animal-Free Recombinant Human EGF (Peprotech), 40 ng/mL Recombinant Human FGF-basic (Peprotech), 200 ng/mL Recombinant Human IGF-I (Peprotech), 10 ng/mL Recombinant Human PDGF-AA (Peprotech) and 10 ng/mL Recombinant Human PDGF-BB (Peprotech).

Both the PDX and the PDO cells were treated with 3.18 uM cisplatin (Cambridge Bioscience). After 1 week the treatment was removed and the cells were kept in normal culture media for 3 weeks. Cells were cryopreserved at each time point (T1 and T3) and before treatment (T0).

After thawing, single cells were washed with PBS and were loaded onto a Chromium Single Cell 3ʹ Chip (10X Genomics) and were run in the Chromium Controller to generate single-cell gel bead-in-emulsions using the 10X genomics 3ʹ Chromium v2.0 platform as per manufacturer’s instructions. Single-cell RNA-seq libraries were prepared according to the manufacturer’s protocol and the library quality was confirmed with a Tapestation High-Sensitivity DNA Kit (Agilent, 5067-5585). Samples were sequenced on an Illumina NovaSeq 6000 according to standard 10X Genomics protocol, aiming for approximately 50,000 reads per cell for gene expression libraries and 1000 reads per cell for CMO/Cellecta barcode libraries.

### Neuroblastoma cell lines treatment for scRNA-seq analysis

The SH-SY5Y and SH-EP cell lines were treated with 3.18 uM cisplatin (Cambridge Bioscience). After 1 week the treatment was removed and the cells were kept in normal culture media for either 3 weeks (SH-SY5Y and SH-EP) or 4 weeks (SK-N-SH). Cells were cryopreserved at each time points (T1 and T3) and before treatment (T0).

After thawing, single cells were washed with PBS and were resuspended with a buffer (Ca++/Mg++ free PBS + 0.04% BSA). Cells were tagged with the CMO oligos (10X Genomics) as per manufacturer’s instructions. Samples were multiplexed in groups of 3 or 4, for a total of 10000 cells/pool.

Pooled single cell suspensions were loaded on a Chromium Single Cell 3ʹ Chip (10X Genomics) and were run in the Chromium Controller to generate single-cell gel bead-in-emulsions using the 10X genomics 3ʹ Chromium v2.0 platform as per manufacturer’s instructions. Single-cell RNA-seq and CMO libraries were prepared according to the manufacturer’s protocol and the library quality was confirmed with a Tapestation High-Sensitivity DNA Kit (Agilent, 5067-5585). Samples were sequenced on an Illumina NovaSeq 6000 according to standard 10X Genomics protocol, aiming for approximately 50,000 reads per cell for gene expression libraries and 1000 reads per cell for CMO libraries.

## Data availability

The raw sequencing data supporting the findings of this study are available in the National Centre for Biotechnology Information - Sequence Read Archive (NCBI - SRA). This can be accessed using the following link: https://dataview.ncbi.nlm.nih.gov/object/PRJNA1071370?reviewer=m1girilboh9soufg9bhoio9d9k. The intermediate processing files are available on Zenodo via the following link: https://zenodo.org/records/10568834.

## Code availability

The codes used for this study are available the GitHub platform. Code to generate mathematical modelling is available at https://github.com/sarthak-sahoo-0710/neuroblastoma_plasticity. Code directory used to process single cell RNA sequencing FASTQ files is available at https://github.com/MikeDMorgan/Genomics. All other codes used for analysis in this manuscript are available in the following GitHub repository: https://github.com/PCEBrunaLab/Roux-et-al.-2024.

## Acknowledgements

We thank Katerina Rekopoulou and Nik Matthews for assistance with sample and library preparation for single-cell RNA-sequencing; Jan Molenaar for providing the patient-derived spheroid lines; Andrea Sottoriva for sharing the Cellecta barcode library and expertise in barcoding the cells as well as Salvatore Milite and Haider Tari for assistance and technical advice in analysis and barcode sequencing processing; Yura Grabovska and Chris Jones for technical expertise in the analysis of scRNA-seq data; Melanie Beckett, Elizabeth R. Tucker, Evon Poon and Karen Barker for helpful discussion and technical advice; Peijie Zhou for help with the MuTrans analysis; Oscar M. Rueda and Maurizio Callari for technical advice regarding WGS and WES analyses. The PDX model was developed within the MAPPYACTS study following informed consent from patients and legal representatives(*41*, *42*). The authors thank all patients and parents/caregivers who participated in the Mappyacts trial and consented to the development of preclinical models. We acknowledge the ICR facilities: Light Microscopy and Confocal, Flow Cytometry, Genomics and Scientific Computing. We acknowledge the RSE Group at The Institute of Cancer Research for providing software enhancements, specifically Rachel Alcraft. Their support was instrumental in achieving the results presented in this paper. A.B. is supported by ICR London. This study was supported by Little Princess Trust Innovation Grant LPTINS2\9 (to A.B and L.C).

## Author contributions

Methodology, C.R., S.H., A.S., E.C., C.E., A.S.S., A.A.A.; experimental design, C.R., S.H., A.S., M.D.M., A.B..; analytical investigation, S.H., E.C., A.S.S., M.D.M., A.B.; writing–original draft, C.R., A.B.; writing–review and editing, C.R., S.H., A.S., E.C., C.E., M.D.M., A.B.; resources, L.C., G.S., B.G.; conceptualisation, A.B.; project administration, A.B.; funding acquisition, L.C., A.B.

## Notes

### Competing Interest Statement

The authors have declared no competing interest.

### Summary of Updates

All results sections have been revised, and the manuscript now includes eight main figures and nineteen supplementary figures to better illustrate the findings. The list of authors has also been updated.

